# Massively parallel assessment of gene regulatory activity at human cortical structure associated variants

**DOI:** 10.1101/2025.02.08.635393

**Authors:** Nana Matoba, Jessica C. McAfee, Oleh Krupa, Jess Bell, Brandon D. Le, Jordan M. Valone, Gregory E. Crawford, Hyejung Won, Jason L. Stein

## Abstract

Genetic association studies have identified hundreds of largely non-coding loci associated with inter-individual differences in the structure of the human cortex, though the specific genetic variants that impact regulatory activity are unknown. We implemented a Massively Parallel Reporter Assay (MPRA) to measure the regulatory activity of 9,092 cortical structure associated DNA variants in human neural progenitor cells during Wnt stimulation and at baseline. We identified 918 variants with regulatory potential from 150 cortical structure associated loci (76% of loci studied), of which >50% showed allelic effects. Wnt stimulation modified regulatory activity at a subset of loci that functioned as condition-dependent enhancers. Regulatory activity in MPRA was largely induced by Alu elements that were hypothesized to contribute to cortical expansion. The regionally specific impact of genetic variants that disrupt motifs is likely mediated through the levels of transcription factor expression during development, further clarifying the molecular mechanisms altering cortical structure.

## Introduction

The expansion of the cerebral cortex in humans, relative to non-human primates, is thought to contribute to our distinctive cognitive and social abilities^1,2^. Genetic association studies conducted in large consortia, including the Enhancing Neuroimaging Genetics through Meta-analysis (ENIGMA), have identified single nucleotide polymorphisms (SNPs) associated with inter-individual variability in human cortical surface area and thickness measured with magnetic resonance imaging, providing a foothold into the molecular mechanisms affecting human cortical size and shape^3^. However, most cortical structure associated variants are found in non-coding regions of the genome without a clearly defined function.

Cortical structure associations are enriched in regulatory regions of the genome active in primary human neural progenitor cells (phNPCs) and colocalize with genetic variants influencing chromatin accessibility - a proxy for transcription factor (TF) binding - and gene expression specifically in NPCs (chromatin accessibility and expression quantitative trait loci; ca/eQTLs)^3–6^. This suggests cortical structure associated variants alter gene regulatory elements by modifying TF binding and gene expression, impacting fate decisions of phNPCs during prenatal development, ultimately leading to differences in cortical size in adulthood. TFs exhibit regionally specific expression patterns^7^, and translocate to the nucleus in response to activation of developmental signaling pathways such as Wnt^8^, which may lead to variants having stronger or weaker effects within specific regions or in stimulated conditions. This hypothesis is supported by cortical structure associations enriched within Wnt responsive regulatory elements and associations found near genes involved in the canonical Wnt pathway^3,6^.

Gene regulatory elements are marked by accessible chromatin or histone post-translational modifications that allow TF binding, and through looping with promoters, can influence gene expression^9–12^. In addition to chromatin loops, enhancer RNAs (eRNAs), transcribed in non-coding genomic regions, initiate looping through complementarity to upstream antisense promoter RNAs (uaRNAs) recruiting polymerase to increase transcription^13,14^. The latter mechanism involves co-opting the expression of Alu elements, a set of transposable elements that are prevalent across the human genome^15^. Alu elements have been theorized to strongly contribute to human cortical evolution because they have increased in prevalence together with brain size along the human lineage^16^.

Due to the correlation of nearby genetic variants (linkage disequilibrium or LD), the specific variant(s) with gene regulatory activity in an associated locus are unknown. We implemented a Massively Parallel Reporter Assay (MPRA), a high-throughput experiment designed to measure the regulatory activity of short DNA sequences using barcoded expression constructs to identify cortical structure associated variants with gene regulatory activity^17,18^. MPRA can measure the regulatory function of a given element (MPRA activity) as well as the influence of an allele within the regulatory element (allelic effect or expression modulating variant [emVar]). Here, we tested the regulatory impact of 9,092 cortical structure associated variants within phNPCs. Considering the role of the Wnt signaling pathway in brain size^19^, we tested whether regulatory potential of variants changes in response to Wnt stimulation. We identified 1,258 elements (7% of elements tested) with regulatory potential from 150 cortical structure associated loci (76% of loci studied). Among these, >50% of cortical structure associated variants showed allelic effects. Regulatory activity in MPRA was largely driven by Alu elements, a small subset of which showed condition-dependent gene regulatory function. Disruption of motifs by emVars highlighted TFs likely involved in human brain development and suggested that the impact of genetic variants on different brain regions is mediated by levels of TF expression in those regions. Overall, characterization of MPRA-validated cortical structure associated variants further clarify molecular mechanisms altering cortical structure.

## Results

### MPRA design and reproducibility

We designed four sets of 150 base pair elements to understand the gene regulatory and allelic effects governing human brain development and structure (**Fig. 1a,b**): (1) *positive controls*: 10 constitutively active promoter regions commonly used to drive transgene expression (min-tk promoter, Ef1-alpha core promoter, CMV promoter) (2) *caQTLs*: 502 chromatin accessibility peaks that contain bi-allelic SNPs replicably associated with their accessibility in a population of phNPCs^4,6^ (**Supplementary Fig. 1**), (3) *ENIGMA GWAS*: 9,092 regions centered on any SNP associated (*P* < 1 x 10^-5^) with inter-individual differences in human cortical structure from 198 genome-wide significant loci^3^ and (4) *negative controls*: 194 scrambled elements from the ENIGMA GWAS. In total, we tested the regulatory activity of 19,500 elements.

**Fig.1.**
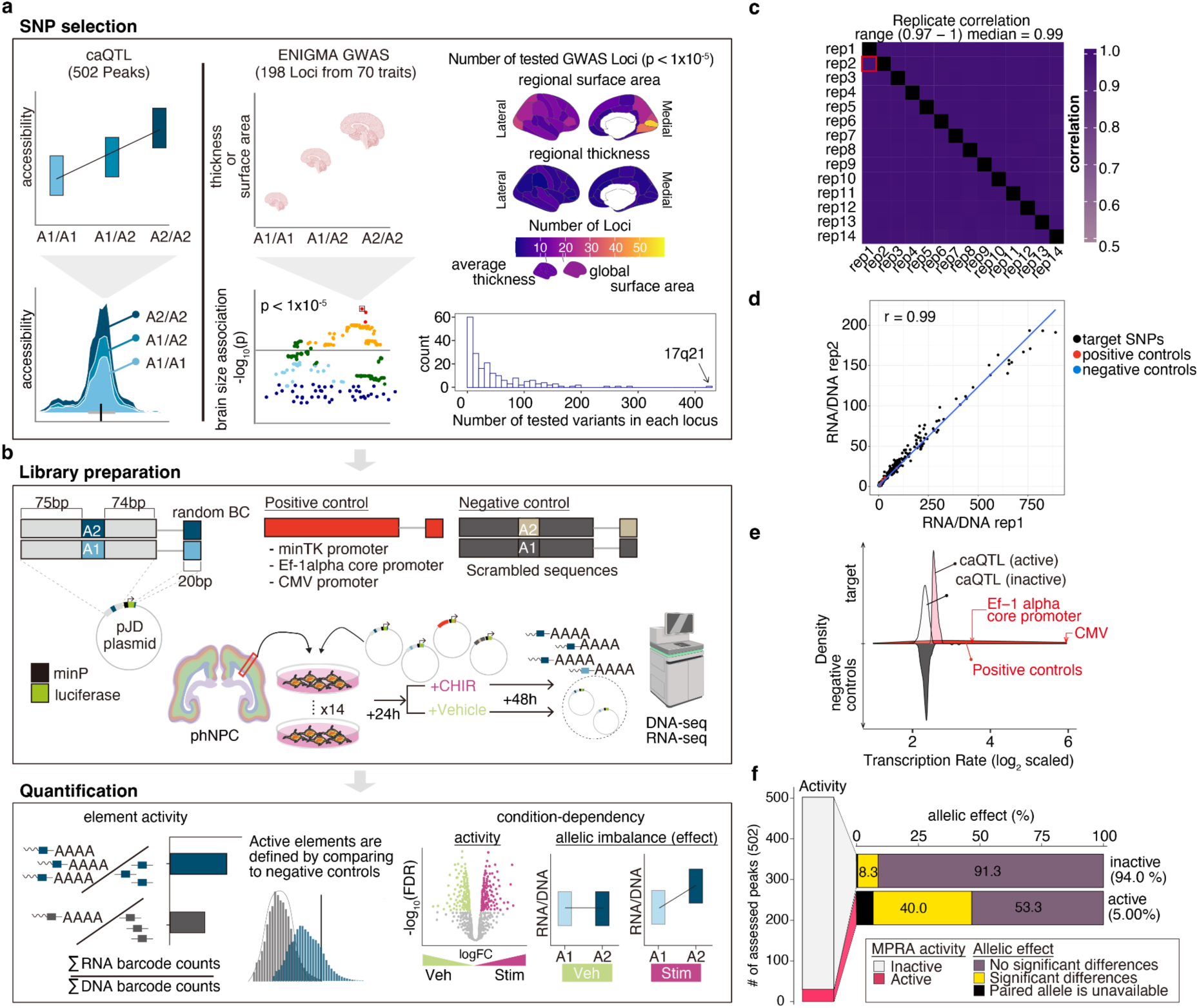
Experimental design and reproducibility. **a,** SNP selection. 502 caQTLs were chosen in chromatin accessible peaks in phNPCs (left). ENIGMA GWAS data contains SNPs associated with surface area or thickness (global and 34 regions of the cortex) (right). Cortical regions are color-coded according to the number of loci included in this study, stratified by surface area and thickness (top) and the number of elements per locus (bottom). **b**, Experimental design. **c**, Replicate correlations of MPRA activity were uniformly strong (median = 0.99). **d**, An example correlation of MPRA activity across two replicates. Each set of MPRA elements is colored along with a line of best fit. **e**, Histograms of MPRA activity, quantified as transcription rate, for different sets of MPRA elements, where positive controls showed higher MPRA activity than negative controls (see also **Supplementary Fig. 4**). For visualization purposes, the distribution of negative controls are shown upside down. **f**, MPRA activity of caQTL elements above the negative controls (MPRA active; FDR < 0.1) was detected in either allele of 37 regions from 502 assessed. Allelic expression imbalances (allelic effects) were observed more frequently in MPRA active elements.

Each element was synthesized and cloned into a plasmid upstream of a minimal promoter to drive expression of luciferase followed by a random 20 bp barcode^20^. We transfected the pooled MPRA library into 16 biological replicates of phNPCs. After 24 hours to allow expression of the MPRA constructs, we activated Wnt signaling using CHIR (CHIR99021, also known as CT99021), a potent GSK3β inhibitor and Wnt activator, or studied baseline gene regulation using a vehicle (DMSO), for 48 hours^3,8,21^. For each biological replicate, we measured the expression of barcodes using RNA-seq and the number of barcodes present in that replicate using DNA-seq. We calculated the MPRA activity of an element as the summed RNA counts divided by the DNA counts^20,22^.

After removing outliers in expression level and filtering poorly represented elements (**Methods**, **Supplementary Fig. 2**), we retained 28 (s.d. = 12.5) barcodes on average for each element across 14 replicates (**Supplementary Fig. 3**). Regulatory activity was strongly correlated between biological replicates, demonstrating the reproducibility of our assay (median Pearson’s correlation = 0.99; **Fig 1c,d**). At least one element from each positive control region showed significantly higher regulatory activity than negative controls (Benjamini-Hochberg (BH)-adjusted *P*-value [FDR] < 0.1; **Fig. 1e, Supplementary Fig. 4**), as expected.

We next evaluated the concordance between caQTLs from our previous work^4,6^ and MPRA conducted in the same cell type - phNPCs (**Supplementary Tables 1,2)**. Though there was limited concordance between MPRA activity or allelic effects in caQTLs and MPRA, as found in previous work^23–27^ (**Supplementary Fig. 5**), the strongest predictor of detecting MPRA allelic effects from caQTLs was MPRA activity (Odds Ratio [O.R] = 8.22; two-sided Fisher’s Exact test *P* = 3.2 x 10^-6^, **Fig. 1f, Supplementary Table 2**). Our subsequent downstream analyses only evaluated allelic effects when at least one allele showed MPRA activity significantly above negative control samples.

### Identifying regulatory elements associated with inter-individual differences in human cortical structure

To identify cortical structure associated regulatory elements, we empirically tested the regulatory activity of genetic variants associated with inter-individual variability in human cortical structure from the ENIGMA GWAS^3^ (**Fig. 1a, Supplementary Table 3a**). Cortical regions varied in the number of detected GWAS significant loci (e.g., 54 loci associated with pericalcarine surface area), and the number of associated SNPs within a locus due to LD patterns (e.g., 428 variants were tested at the large chromosome 17q21 inversion region^28^), influencing the number of elements tested in our MPRA (**Fig. 1a, Supplementary Table 3a**). The vast majority (88.4%) of variants tested in the MPRA were located in introns or intergenic regions, with presumed gene regulatory function. Twenty percent of variants were associated with both cortical surface area and thickness, suggesting these variants have pleiotropic effects (**Supplementary Fig. 6a, Supplementary Table 3a**).

Among 17,837 elements (9,052 variants) in 198 loci, we observed 988 elements with MPRA activity significantly higher than the negative controls in the vehicle condition (FDR < 0.1, **Fig. 2a,b, Supplementary Fig. 6, Supplementary Tables 1,3a**). While only 6% of tested elements showed MPRA activity, 70% of loci contained at least one element with detectable regulatory activity. This suggests that most cortical structure associated loci have at least one SNP with detectable regulatory activity in phNPCs using MPRA.

**Fig. 2.**
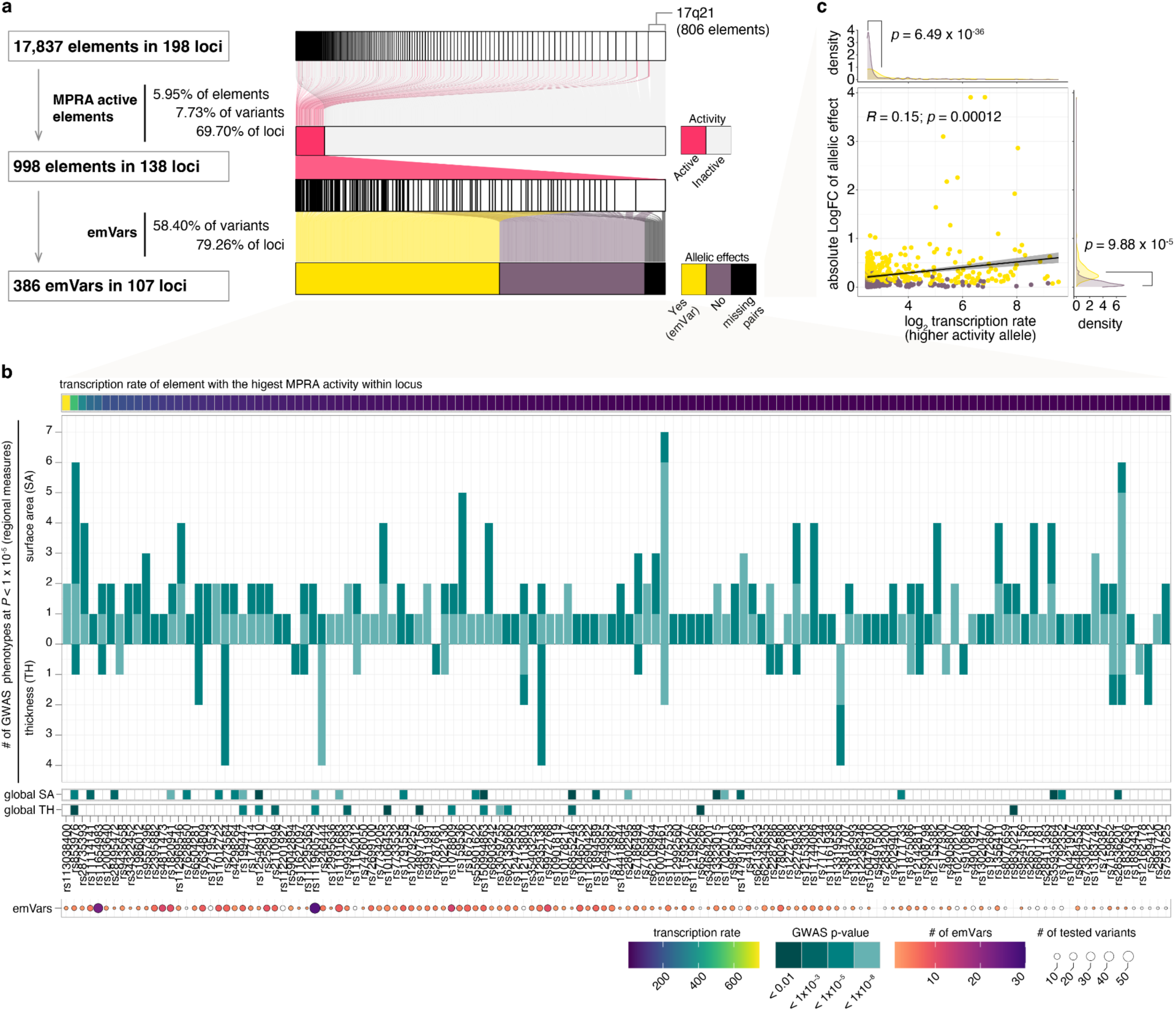
MPRA regulatory activity and allelic effects at cortical structure associated loci. **a,** Summary of MPRA elements at cortical structure associated loci with MPRA activity and allelic effects. Sankey diagram illustrates the composition of elements across tested loci (top), colored by MPRA activity (middle) or emVars (bottom). emVars were tested only for variants where at least one allele showed MPRA activity. **b,** MPRA active cortical structure associated loci ordered by MPRA activity. *Top row:* the level of MPRA activity of the allele with highest activity within a locus. *Middle row:* a barplot showing the number of cortical regions (among 34) that were associated to a locus containing an MPRA active element colored by GWAS significance level, followed by GWAS *P-value* of global surface area and thickness. *Bottom row:* the number of emVars at the locus. **c**, MPRA transcription level of the higher activity allele is associated with the absolute value of allelic difference (logFC). The line was fitted using linear regression and shadows indicate 95% confidence interval. Pearson correlation coefficient (R) and its *P*-value are shown.

More MPRA active elements were discovered at surface area associated loci than at thickness associated loci, likely due to the larger number of elements tested for surface area (**Supplementary Fig. 6**). MPRA active elements generally had regionally limited effects, being significantly associated with only one or two cortical regions (GWAS *P* < 1×10^-5^) rather than having broad impacts across the entire cortex (**Fig. 2b, Supplementary** Fig. 6**, Supplementary Table 3a**).

We next evaluated allelic effects at cortical structure associated loci that showed MPRA activity when both alleles were present. We found 386 variants with significant allelic effects (79% of loci, 58% of variants tested; FDR < 0.1; **Fig. 2b, Supplementary Tables 1,3b**), indicating that most cortical structure associated loci have at least one genetic variant capable of altering transcription in progenitor cells during brain development. Consistent with the caQTL dataset, emVars exhibited higher expression levels than non-emVars, further supporting the relationship between MPRA expression strength and allelic effects (**Fig. 2c**).

### Cortical structure associated MPRA active elements are enriched in promoter regions of Alus

We next aimed to identify genetic features associated with MPRA active elements linked to cortical structure. We found that MPRA active elements were enriched in short interspersed nuclear elements (SINEs; 56.7% in active, 11.6% in inactive elements), consistent with previous MPRA experiments in other cell types^27,29^ and phNPCs^20^ (**Fig. 3a**). SINEs are a type of the retrotransposons comprising a set of DNA elements that express themselves and re-integrate into new positions, resulting in a large number of repeated copies found throughout the genome. Retrotransposons have been hypothesized to contribute to the rapid evolution of both gene regulation and human brain structure^16,30^. Among subfamilies of SINEs^31^, MPRA active elements were specifically enriched in forward oriented Alu elements^32^ (**Fig. 3b-d**), which aligned with our oligo design that used only forward sequences.

**Fig. 3.**
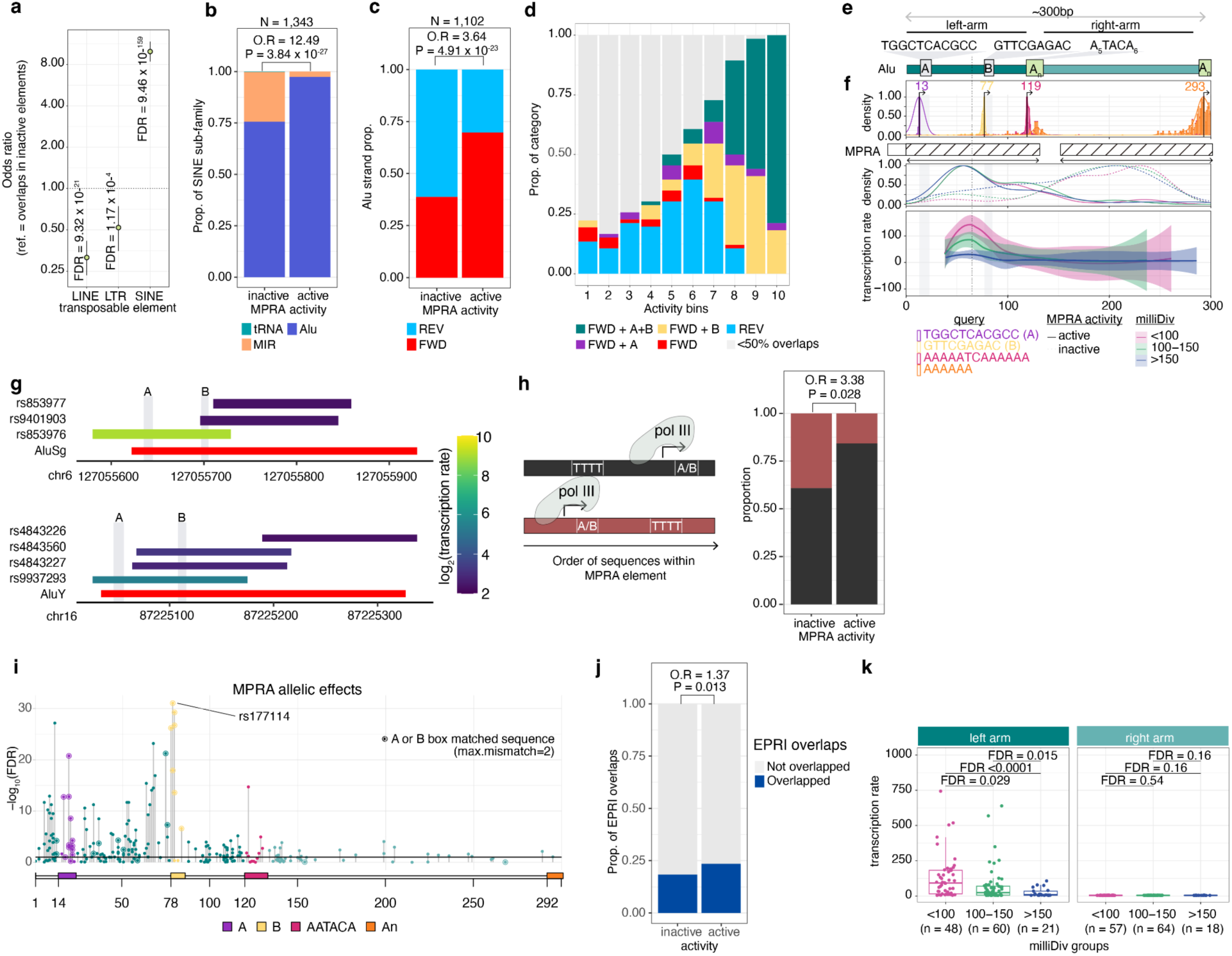
MPRA activity of cortical structure associated loci is driven by Pol III promoter elements within Alus. **a,** Enrichment of MPRA active elements across classes of transposable elements, **b**, SINE-subfamily elements, and **c,** Alu element orientation, showing that forward Alu elements are major drivers of MPRA activity. **d,** MPRA active elements binned by level of activity were colored based on their overlap to Alu elements in different orientations (FWD/REV) or containing A and B box Pol III promoter elements. **e,** Schematic structure of a typical Alu-element which contains a left arm serving as Pol III promoter containing A and B boxes, followed by a right arm terminating with a polyA repeat. **f,** Frequency of MPRA elements containing sequence corresponding to the positions of observed consensus elements in the Alu element, including A and B boxes (top). Density of MPRA elements stratified by MPRA activity group across position (middle). Activity level of MPRA active elements graphed according to their position within the Alu element, stratified and colored using base mismatches in parts per thousand (milliDiv) (bottom). **g,** Example MPRA elements demonstrating that higher MPRA activity (color) is observed when the elements contain both A and B boxes. **h,** Elements where A/B matched sequences occur before the Pol III termination sequence (TTTT) are more often inactive. **i,** Significance of MPRA allelic effects mapped within the Alu element show that genetic variation near A/B boxes have strong effects on MPRA activity. **j,** Overrepresentation of MPRA active elements within enhancer RNAs or upstream antisense RNAs assessed in neural progenitor cells using RIC-seq^13^. **k,** Younger elements with lower milliDiv show higher activity than older elements, specifically when found in the left arm of the Alu element. Statistical significance of enrichment was assessed by two-sided Fisher’s exact test. *P-*values were corrected for multiple comparisons using the Benjamini–Hochberg method. FWD: Forward, REV: Reverse, Veh: vehicle, O.R: odds ratio, EPRI: enhancer-promoter RNA interaction.

A typical Alu element contains two consensus sequences in its left arm that function as RNA polymerase III (Pol III) promoters, the A-Box (TGGCTCACGCC) and the B-Box (GTTCGAGAC)^33,34^ (**Fig. 3e**). We estimated the relative position of MPRA elements within the Alu sequence, and observed that the most active MPRA elements were located in the left arm of the Alu element and contained the A or B boxes (**Fig. 3d-g**). Presence of both A and B boxes resulted in the highest MPRA activity (**Fig. 3d-g**). One of the scrambled sequences (negative control) unintentionally contained a B-box like sequence (GTTCGAGC) and had strong MPRA activity, further supporting that these Alu based sequences drive MPRA activity (**Supplementary Fig. 7**). We also found reduced MPRA activity when elements containing A or B boxes were followed by a TTTT (T4), the termination signal for Pol III (**Fig. 3h**). Subsequently, we explored allelic effects within MPRA active elements that contained Alu sequences. We found that emVars were more often found in Alu elements compared to MPRA active variants without allelic effects (O.R = 1.56, P = 0.007). Notably, the variants with the most significant allelic effects on transcription were located within the left arm of the Alu element and overlap or were close to A or B boxes, further supporting the critical regulatory role of these regions (**Fig. 3i**).

Strong Alu-element driven transcription may be an artifact of non-chromatinized sequences in the episomal MPRA library, that are normally silenced in the endogenous genome. We found that MPRA active elements were enriched with eRNAs and uaRNAs in phNPCs^13^ compared to MPRA inactive elements (**Fig. 3j**), suggesting that at least a subset of MPRA active elements exhibit endogenous genome activity.

The age of Alu elements can be estimated based on their accumulated mutations over time, with younger Alu elements having lower sequence divergence to ancestral sequences than older Alu elements. The level of sequence divergence can be measured using mismatches per thousand bases to consensus sequences (milliDiv)^35^. We found that younger cortical structure associated Alu elements (<100 milliDiv) showed higher activity than older elements (>150 milliDiv) (**Fig. 3f,k**), consistent with previous findings^32^. These results suggest that more recent insertion of Alu elements created novel gene regulatory regions impacting neurodevelopmental genes that influenced cortical structure along the human lineage.

Overall, these results demonstrate that MPRA activity of cortical structure associated sequences in neural progenitors are strongly driven by Alu elements, especially younger Alu, that enhance Pol III based transcription.

### Transcription factors contributing to regulatory function at cortical structure associated elements

Since TF binding is known to regulate gene expression, we explored whether certain TF binding motifs contribute to MPRA activity. Elements with MPRA activity had more complex sequences as evaluated by Shannon entropy^36^ (*p* = 4.58 x 10^-24^ using a t-test) and were more likely to contain a TF binding motif (**Fig. 4a**), as has been found in previous MPRA studies^22,25,37^. Specifically, three separate clusters of similar TF binding motifs (TFBMs) – ZNF135/460, MEF2A-D, and Zfx – were enriched in human cortical structure associated MPRA active elements in phNPCs (**Fig. 4b**). ZNF460 TFBMs were previously found to be enriched in great ape-specific structural variants and hypothesized to increase brain size along the human lineage^38^. Conditional knockout of *Mef2c* in mice was shown to reduce whole brain size^39^. *MEF2C* is also involved in neuronal differentiation, and genetic variation within or near this gene creates risk for psychiatric disorders^22,40^. Alu elements contain sequences similar to ZNF135, ZNF460, and MEF2A binding motifs which exhibit human-specific activity and may be driving this observed overlap^41,42^.

**Fig. 4.**
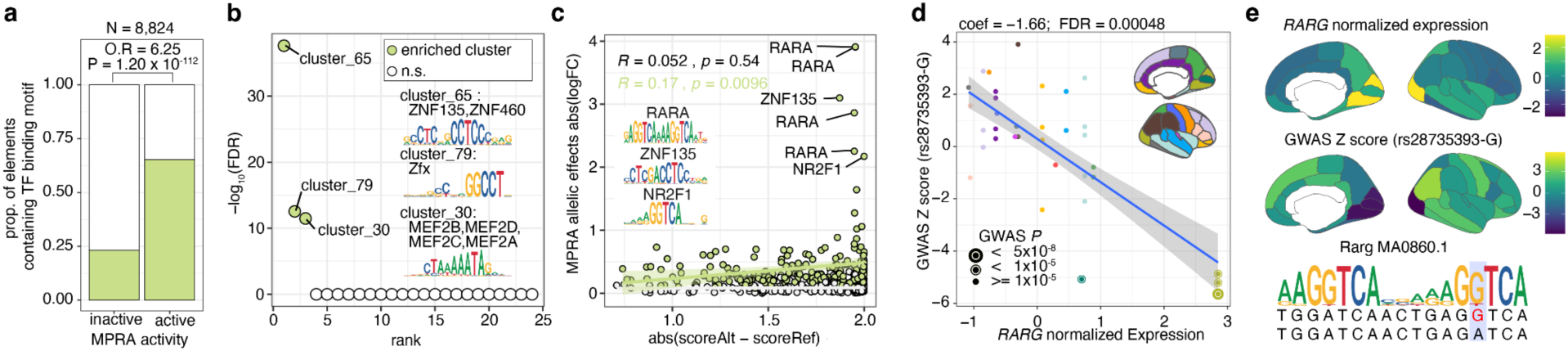
TFs enriched in cortical structure associated regulatory elements and underlying regional specificity. **a,** Proportion of elements containing TF binding motifs stratified by MPRA activity. **b,** Enriched TF binding clusters. Statistical significance of enrichment in **a** and **b** were assessed by a two-sided Fisher’s exact test. **c**, The level of TF motif disruption (x-axis) is related to MPRA allelic differences (y-axis). *P*-values were estimated by Pearson’s correlation test. **d-e,** TF expression levels during prenatal development across brain regions (colored) were associated with the impact of variant on cortical structure. Each dot represents DK-atlas brain area colored by macro-scale brain area (**d**) (**Supplementary Fig. 8**). **e**, Brain areas are colored by scaled expression values (top) and GWAS Z-scores (middle). Seqlogo showing G allele matches the Rarg binding motif site (bottom). In **c** and **d**, the lines were fitted using linear regression and shadows indicate 95% confidence interval. *P-*values were corrected for multiple comparisons using the Benjamini– Hochberg method (**b,d**). O.R: odds ratio.

Previous studies have shown that the level of motif disruption is associated with MPRA allelic effects^22,37^, so we evaluated this relationship in our data. We found that emVars strongly correlated with the predicted level of motif disruption, whereas elements without allelic effects had no detectable relationship with motif disruption (**Fig. 4c**). Our results suggest that TF binding affinity impacts the strength of allelic effects on transcription observed in MPRA. Notably, disruption of RARA, ZNF135, and NR2F1 motifs led to the largest impacts on MPRA activity. It was reported that the consensus sequences of evolutionary recent Alu subfamiles contain binding sites for retinoic acid receptors^43^, consistent with our finding of stronger MPRA activity within younger Alu elements (**Fig. 3f,k**). This indicates TFs that may shape the human cortex during development.

### Spatial relationships between TF expression and allelic effects

Genetic variants often are associated with differences in a limited number of cortical regions rather than having global effects across all cortical regions, despite the fact that the variants are constant in all cells (**Fig. 2b**). We hypothesized that the molecular mechanism underlying regionally specific cortical structure effects is that regions more affected by a given variant would also have higher expression of TFs disrupted by that variant, similar to findings from previous studies^44,45^.

To test this hypothesis, we compared predicted TF binding motifs disrupted by emVars to corresponding TF expression levels across 16 macroscale brain regions (**Supplementary Fig. 8**) from 8 to 24 post-conception weeks (pcw) from the BrainSpan Atlas^46^. We then analyzed the spatial correlation between TF expression and the impact of the variant on cortical structure^3^. We found a significant correlation between GWAS effect sizes for 7 cortical structure associated SNPs and regional TF expression (**Supplementary Tables 4,5**). For example, elevated *RARG* expression was associated with a greater reduction in surface area in the presence of rs28735393-G, which matches the Rarg binding motif (**Fig. 4d,e**). Our results suggest that TF expression levels, in combination with genetic variants disrupting TF binding, modulate the impact of genetic variants on cortical structure.

### Wnt activity modulating MPRA activity

Wnt stimulation alters gene regulation to modulate proliferation and differentiation of NPCs shaping brain structure^6,8,19^. To evaluate the condition-dependent activity of cortical structure associated variants, we stimulated the Wnt pathway in phNPCs using CHIR (**Methods**). We detected a similar number of MPRA active elements (927 among 17,848 elements; 6.94% of tested variants and 67.17% of tested loci) in the stimulated condition (**Fig. 5a,b**, **Supplementary** Fig. 6, **Supplementary Tables 1,3c)**. The same TF clusters were enriched in MPRA active elements under both the vehicle and Wnt stimulated conditions, likely due to 657 active elements being detected in both conditions (**Supplementary Fig. 9**). While we detected 270 elements that were active only in the Wnt stimulated condition (**Fig. 5b**), many of these regions are likely just above or below the threshold cutoff. To more stringently evaluate condition dependency, we further identified MPRA active elements with significantly different levels of activity across conditions. We found 32 elements with higher activity due to Wnt stimulation and 7 elements with higher activity in the vehicle condition, demonstrating Wnt-dependent gene regulation of cortical structure associated elements (**Fig. 5b,c, Supplementary Table 3d**). MPRA active elements with higher activity in simulated conditions more often overlapped with SINEs compared to elements whose activity did not significantly differ between conditions (**Fig. 5d**).

**Fig. 5.**
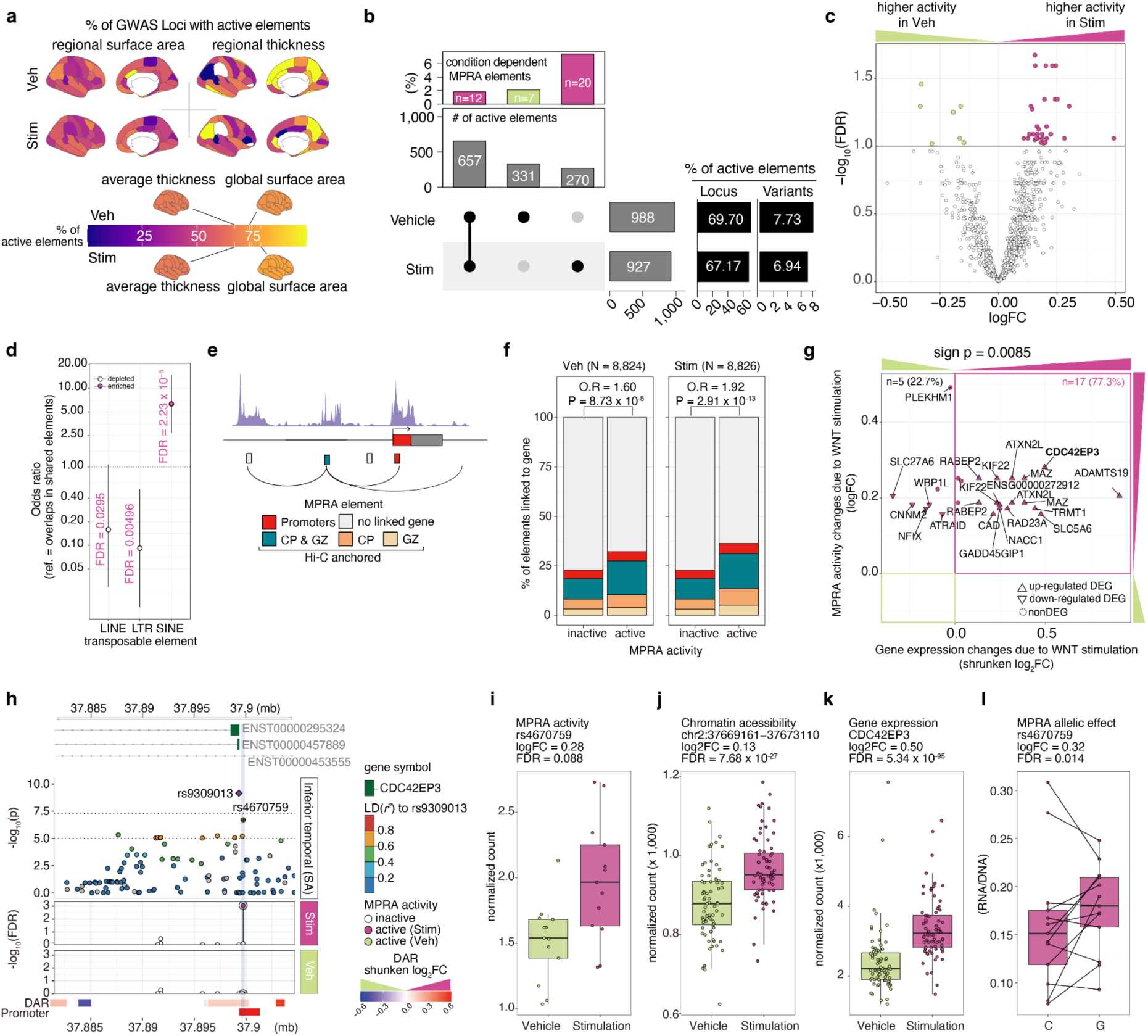
Regulatory elements associated with cortical structure show condition-dependent function. **a,** The percentage of MPRA active elements across brain regions stratified by condition (top: Vehicle, bottom: Wnt Stimulation via CHIR) and measurement (left: surface area, right: thickness). **b,** The number of MPRA active elements in each condition (bottom). The top panel shows the percentage of condition-dependent MPRA elements. **c,** Volcano plot showing condition-dependent MPRA activity in Wnt stimulation vs vehicle. **d,** Enrichment of TEs within elements exhibiting higher activity during Wnt stimulation compared to elements without detectable differences in MPRA activity across stimulation conditions. **e,** Schematic example linking MPRA element to genes at their promoter region or using chromatin interaction (Hi-C) for distal elements. **f,** Percentage of MPRA elements linked to genes stratified by MPRA activity and condition. Active elements were more likely to be linked to a gene. **g,** Scatter plot showing gene expression changes vs MPRA activity changes due to Wnt stimulation. A sign test was used to determine if condition-dependent MPRA activity occurred in the same direction as condition-dependent gene expression more than expected by chance. **h**, *CDC42EP3* locus highlighting a condition-dependent MPRA element (rs4670759). Top: - log10(p-value) from GWAS of inferior temporal cortex surface area. Each dot represents variants in the locus colored by LD (*r*^2^) to the index SNP (rs9309013, purple diamond). Middle: -log_10_(FDR) of MPRA activity in stimulation or vehicle conditions. Bottom: Differential chromatin accessible peaks colored by shrunken log2FC, followed by a promoter region. **i-l,** Boxplots displaying differences between conditions in multiple measurements: MPRA activity (**i**), chromatin accessibility (**j**), gene expression (**k**) in phNPCs. **l,** MPRA allelic effects of rs4670759 in stimulated condition. Dots indicate replicates/donors in **i**-**l**. Same replicate was connected by line in **l**.; *n* of replicates/donors: *n*_MPRA_ = 14, *n*_chromatin_accessibility_ = 72, *n*_gene_expression_ = 74. *P*-values were corrected for multiple comparisons using the Benjamini-Hochberg procedure. Stim: Stimulation, Veh: Vehicle, CP: cortical plate of the developing cortex; GZ: germinal zone of the developing cortex.

### Condition-dependent regulation of gene expression

In order to determine downstream impacts of condition-specific MPRA active elements, we mapped them to putative target genes. MPRA active elements in gene promoters were directly assigned to their target genes, while the rest were mapped to the target genes via chromatin interaction profiles in the developing brain^10^. We found that MPRA active elements were more likely to be linked to a gene than MPRA inactive elements (**Fig. 5e, f**). Interestingly, elements with increased MPRA activity under Wnt stimulation were linked to genes with higher expression under the same condition^6^, suggesting these elements function as condition-dependent enhancers^6^ (**Fig. 5g**). As an example, rs4670759, a SNP associated with the surface area of the inferior temporal cortex, showed higher MPRA activity (**Fig. 5h**) and increased chromatin accessibility (**Fig. 5i,j**) under Wnt stimulation. This variant is located in the promoter of *CDC42EP3* (**Fig. 5h**), which is involved in actin cytoskeleton formation^47^ and whose expression is increased under Wnt stimulation^6^ (**Fig. 5k**). We note this element is an emVar in the stimulated condition, though we were unable to test for allelic effects in the vehicle condition because one of the alleles was missing in the QCed dataset (**Fig. 5l**). In summary, we present evidence that condition-dependent MPRA activity of cortical structure associated SNPs results in condition-dependent effects on gene expression.

### Condition-dependent allelic effects at cortical structure associated loci

Many variants (*n* = 183) showed MPRA allelic effects regardless of stimulation, but more variants were identified as emVars in only one condition (*n*_emVars_stimulation only_ = 150, *n*_emVars_vehicle only_ = 203), suggesting Wnt stimulation can drive condition-specific variant function (**Fig. 6a,b, Supplementary Table 3b,e**). By using an interaction test, we detected 12 variants with allelic effects that differ significantly across conditions (condition-dependent emVars) (**Fig. 6b**, **Supplementary Table 3f**). Of those, none disrupted TCF/LEF motifs which are TFs downstream of canonical Wnt pathway activation^49^. However, TFs disrupted by these stimulus-interacting SNPs were differentially expressed in phNPCs following Wnt stimulation, suggesting a mechanism by which cortical structure associated variants exert condition-specific allelic effects (**Fig. 6c**).

**Fig. 6.**
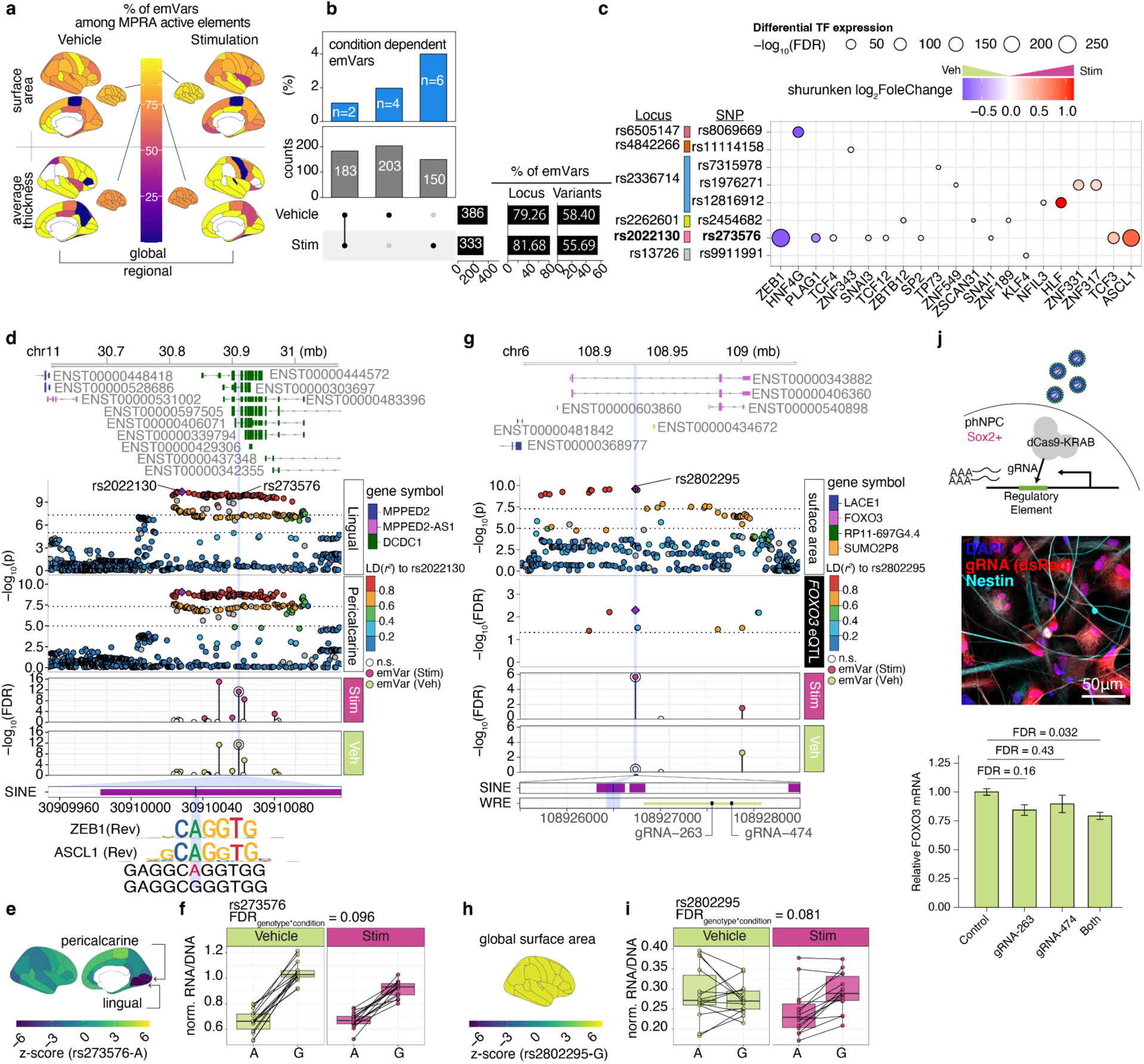
Condition-dependent allelic effects in cortical structure associated regulatory elements. **a,** Percentage of emVar across brain regions stratified by condition (left = vehicle, right = stimulation) and measurements (surface area (top), thickness (bottom)). **b,** Number of emVars in each condition (bottom). The percentage of condition-dependent allelic effects (top). **c,** Disrupted TFs and their Wnt induced expression at condition-dependent allelic MPRA elements. The size of the dots indicates the significance of differential expression due to Wnt stimulation colored by shrunken log_2_FoldChange. **d**, *DCDC1* locus highlighting condition-dependent allelic MPRA element (rs273576). Top; -log_10_(pvalue) from GWAS of Lingual and Pericalcarine surface area. Each dot represents variants in the locus colored by LD (*r*^2^) to the index SNP (rs2022130, purple diamond). Middle: -log_10_(FDR) from MPRA activity in stimulation and vehicle conditions. Bottom: transposable elements and TFs disrupted by rs273576. **e**, Z score of GWAS regional surface area. **f,** Boxplot shows normalized RNA/DNA ratio of MPRA element (rs273576) stratified genotype and conditions. **g**, *FOXO3* locus highlighting condition-dependent allelic MPRA element (rs2802295) (left). Top: -log_10_(pvalue) from GWAS of global surface area and -log_10_(FDR) from *FOXO3* eQTL^48^ (FDR < 0.05 are shown). Each dot represents variants in the locus colored by LD (*r*^2^) to the index SNP (rs2802295, purple diamond). Middle: -log_10_(FDR) from MPRA allelic effects in stimulation and vehicle conditions. Bottom: transposable elements disrupted by rs2802295 and Wnt responsive element (WRE) and the position of the designed gRNAs for the CRISPRi experiment. **h**, Z score of GWAS global surface area. **i**, Boxplot shows normalized RNA/DNA ratio of MPRA element (rs2802295) stratified genotype and conditions. **j**, Experimental design of CRISPRi (top), immunofluorescence image showing expression of gRNA constructs in phNPCs (middle) and relative *FOXO3* expression as measured by qPCR (2 sample t-test with Holm post-hoc correction, *n* = 6 unique donors, mean+/- standard error are shown) (bottom).

An interesting example of a condition-dependent emVar is rs273576 at the intron of *DCDC1*, where Wnt activation diminished the allelic effects detected in the MPRA (**Fig. 6c-f**). The A allele is associated with decreased lingual and pericalcarine surface areas and more closely matches binding sites for ZEB1 and ASCL1, TFs differentially expressed in response to Wnt stimulation^6^ (**Fig. 6c,d**). This variant is an eQTL for *MPPED2* (300 kb downstream) in multiple tissues in GTEx, though not detected in brain or phNPCs.

Another example is rs2802295 within an intron of *FOXO3* (**Fig. 6g**). The G allele is associated with increased global surface area and decreased expression levels of *FOXO3* in the adult brain^48^ (**Fig. 6g**). *FOXO3* gene expression was also downregulated under Wnt stimulation^6^. We detected an allelic effect at this variant under Wnt stimulation that was blunted in vehicle (**Fig. 6i**). This variant is proximal to a Wnt responsive element, where chromatin is less accessible due to Wnt stimulation. To validate the effects of this non-coding region on *FOXO3* gene expression, we performed CRISPR interference (CRISPRi) by designing two gRNAs to target dCas9-KRAB nearby this region (∼1kb from rs2802295) (**Fig. 6j**). We found that silencing of this element by dCas9-KRAB was associated with decreased expression of *FOXO3*, experimentally linking a cortical surface area associated emVar to its gene of action. Using MPRA, CRISPRi validation, and QTLs, we suggest a likely causal variant, gene, and context associated with differences in human cortical structure.

In summary, we identified emVars associated with cortical structure. A subset of these variants showed differential regulatory activity in response to Wnt stimulation, suggesting that signaling pathways critical for brain development may influence the regulatory potential of the genetic variants. These findings reveal how genetic variants shape cortical structure associated differences in the human brain.

## Discussion

Here, we assessed the gene regulatory potential of cortical structure associated variants in phNPCs. We found that most cortical structure associated loci contained elements with regulatory activity in phNPCs, and that cortical structure associated genetic variants influence the level of that regulatory activity. Previous QTL studies using endogenous variation in populations of brain tissue or phNPCs have also found that genetic variants exert effects during neural development to influence adult cortical structure traits^4,6,50,51^. Our MPRA results provide further evidence to this effect using experimental manipulation of alleles, ruling out confounding effects from other variants that exist within an individual.

Because experimental manipulation allows the separation of alleles in high LD, specific genetic variants can be nominated that alter gene regulatory function. Those variants that disrupt TF binding motifs can be used to suggest TFs that lead to changes in cortical structure. Here, we found that TFs like RARA, RARG, ZNF135, and NR2F1 are critical for determining inter-individual differences in human cortical structure. Though previous work on these TFs has been conducted in model systems like mice^52–54^, our findings demonstrate that the TFs impact human brain development and structure. We also propose a mechanism for region-specific effects detected in cortical structure genetic association studies. The regional effects of variants correspond to the expression levels of TFs that bind to them.

Wnt signaling has been well known in non-human model systems and in vitro human model systems to impact brain development^19^. Therefore, we reasoned that the regulatory effect of cortical structure associated variants may vary in response to Wnt stimulation. We find that genetic variants exhibit Wnt stimulation-dependent gene regulatory function (**Fig. 5**), which we have also demonstrated in the endogenous genome using chromatin accessibility and gene expression^6^. Notably, genetic effects can be either accentuated or blunted due to Wnt stimulation. This suggests that multiple cell types and stimulation states will need to be studied to fully elucidate gene regulatory effects impacting cortical structure.

Alu elements are primate specific and have increased in prevalence in the genome coincident with brain size increases along the human lineage^16^. Alu elements transcribe themselves through highjacking proteins derived from other retrotransposable elements and insert themselves into other regions of the genome, providing a mechanism for more rapid gene regulatory evolution than the accumulation of single base pair mutations^15^. Here, we provide evidence that cortical structure associated variants within Alu elements strongly influence transcription in phNPCs. This suggests that Alu elements have been critical for creating gene regulatory elements that harbor RARA, RARG, ZNF135, and NR2F1 motifs, which influence human cortical structure^41,42^. Over time, these Alu elements have likely accumulated genetic variation that alters their function, impacts gene expression during development, and contributes to inter-individual differences in cortical structure.

We found that Pol III promoter regions supplied by Alu elements are strong drivers of MPRA activity. Pol III is generally associated with transcription of short sequences, less than 500 bp in length. Interestingly, the barcode element is contained within the 3’ UTR of the luciferase gene which is ∼1.8 kb from the MPRA element, suggesting luciferase and the barcode are likely transcribed by Pol II rather than Pol III. Given this evidence, we propose a mechanism whereby Pol III transcribes the MPRA element, effectively forming an eRNA that enhances Pol II recruitment, leading to higher expression of the luciferase gene and the barcode contained within. This mechanism has in fact been observed in neural cells^14^.

We implemented MPRA using transfection of episomal plasmids, so our results do not recapitulate endogenous genome chromatinization, one of the major mechanisms underlying gene regulation. Likely because of this, MPRA active elements had limited overlap with chromatin accessibility peaks and MPRA allelic effects had limited overlap with caQTLs that were discovered and replicated in the cell type (**Supplementary Fig. 1,5**). Nevertheless, a subset of MPRA active elements did show overlap with eRNAs or uaRNAs active in phNPCs. Also, condition-dependent MPRA elements showed evidence of mapping to genes with condition-dependent gene expression. This provides evidence that MPRA active elements do have at least partial overlap with activity in the endogenous genome. However, we cannot rule out that MPRA activity from some elements is only due to episomal activity and gene regulatory activity from individual loci would not be recapitulated within the endogenous genome^55^. Future studies generating specific base pair edits in the endogenous genome would be able to disentangle these effects.

## Methods

### Ethics statement for human tissue-derived cell-lines

We acquired human fetal brain tissue following voluntary termination of pregnancy according to IRB regulations at UCLA through the Gene and Cell Therapy Core after consent of the mother for use of tissue in research. No compensation was provided to participants.

### SNP Selection

Several variant sets were generated for inclusion in the MPRA. (1) *positive controls* - Known promoter or enhancer sequences that are commonly used to constitutively drive gene expression (minTk promoter, Ef1-alpha promoter, CMV promoter) were included. If the sequences were >150 bp, arbitrary 150 bp subsets were used. (2) *caQTLs* - SNPs within chromatin accessibility peaks in primary human neural progenitors were selected for inclusion^4,6^. A 150 bp sequence surrounding the variant from the human reference genome was included. Both alleles were synthesized. Indels were removed. (3) *ENIGMA GWAS* - SNPs associated with human cortical surface area or thickness were selected from a large GWAS of cortical surface area and thickness^3^. Both alleles from all variants with *P* < 1×10^-5^ in the association to a cortical structure phenotype and *r*^2^ > 0.6 to the identified index SNPs were included. A 150 bp sequence surrounding the variant from the human reference genome was included. (4) *negative controls* - Negative controls were generated by randomly scrambling the cortical structure associated variant sequences for a subset of those variants. Each allele at the SNP of the unscrambled sequence was included for analysis. Both alleles were removed when the MPRA library sequence contained a restriction enzyme site used for cloning.

### Creating variant oligo library

The 150 bp library oligos mentioned above were synthesized by Agilent and amplified via PCR using the NEBNext 2X Q5 Hifi HS Mastermix (NEB, cat#M0453S) with primers MPRA-chipprimer-R and MPRA-chipprimer-F. Primer information is available in **Supplementary Table 6.**

We then added random 20bp barcodes and restriction sequences (MluI and SpeI) to each variant as previously described^22^ using a pair of primers: MPRA-BC_Primer_R and MPRA-chipprimer-F. Following library preparation, the sample was digested with SpeI-HF (NEB, cat#R3133S) and MluI-HF (NEB, cat#R3198S) for 1 hour at 37°C. This step was immediately followed by a 1-hour rSAP treatment (NEB, cat#M0371S) at 37°C, and then a 5-minute heat inactivation at 65°C. Finally, the digested library was purified using the Zymo Clean and Concentrator Kit-5 (Zymo, cat#D4033).

### Creating MPRA Plasmid Library

The plasmid pJD_Donor_eGP2AP_RC was obtained from Addgene (Addgene, cat#133784). The plasmid was digested with restriction enzymes SpeI-HF and MluI-HF for 1 hour at 37°C, then was heat inactivated for 20 minutes at 80°C, brought to room temperature, followed by rSAP treatment for 1 hour at 37°C, and heat inactivation for 5 minutes at 65°C. The digested plasmid was then gel extracted from 1% agarose gel using Zymo Gel DNA Recovery kit (Zymo, cat#D4007). The digested plasmid and oligo library was ligated together using T7 DNA ligase (NEB, cat#M0318S) at room temperature for 30 minutes followed by clean up via Zymo clean and concentrator-5 kit. The ligation product was transformed into NEB 5-alpha Electrocompetent E. coli (NEB, #C3020K), grown for 7 hours at 30°C in 4L of LB, then maxi prepped using ZymoPURE II Plasmid Maxiprep Kit (zymo, cat#D4203).

### Barcode Mapping

The barcodes (BC)s were mapped to the variants in a method previously described^22^. Briefly, the barcode and variant region was amplified off the plasmid with primers that contain Illumina P5 and P7 adapters (*Bcmap_P5_AAV_R* and *Bcmap_P7_AAV_F_index*) with NEBNext 2X Q5 Hifi HS Mastermix. The PCR product was purified and sequenced using the NovaSeq 6000 SP (2×250 bp) with custom sequencing primers BCmap_R1Seq_AAV_R, BC_Map_R1seq, and BCmap_R2Seq_AAV_F (**Supplementary Table 7**). The code used to map the barcodes to variants can be found in this github repository: https://github.com/thewonlab/schizophrenia-MPRA. There was a mean of 113 unique barcodes per variant prior to transfection.

### Adding in minimal promoter and Luciferase

The luciferase and minimal promoter sequences were obtained from the pGL4.23 plasmid (Promega, cat#E841A). The pGL4.23 plasmid was enzymatically digested with KpnI-HF (NEB, cat#R3142S) and XbaI (NEB, cat#R0145S) at 37°C for 3 hours, followed by heat inactivation at 65°C for 10 minutes. The digestion was cleaned up via Zymo clean and concentrator-5 kit, and then gel extracted using Zymo Gel DNA Recovery kit, yielding a band size around 1.780Kb. The plasmid containing the library and barcodes was digested first with KpnI-HF and rSAP at 37°C for 3 hours, then 65°C for 10 minutes for heat inactivation, cleaned up using Zymo clean and concentrator-5 kit, and then gel extracted using Zymo Gel DNA Recovery kit. Then the product was digested with XbaI and rSAP 37°C for 2 hours, then 65°C for 10 minutes for heat inactivation, and cleaned up using Zymo clean and concentrator-5 kit. The digested plasmid containing the library was ligated with the fragment carrying the minimal promoter and luciferase sequence using T7 DNA ligase at room temperature for 30 minutes. The resulting construct was then purified with the Zymo Clean and Concentrator-5 kit, yielding the final library plasmid, PJD_lib_minP_Luc. PJD_lib_minP_Luc was transformed via heat shock into Library Efficiency DH5α Competent Cells (Invitrogen, cat#18263012) and grown in kanamycin LB overnight at 30°C. The PJD_lib_minP_Luc plasmid was maxi prepped with ZymoPURE II Plasmid Maxiprep Kit.

### Transfecting PJD_lib_minP_Luc into phNPCs

Acquisition of phNPCs and cell culture conditions were previously described^50^. Donor number 88 was used for these experiments and is biologically male. 6-well plates were coated with Poly-L-Ornithine (10 μg/ml; Sigma-Aldrich, cat#P3655-100MG) and Laminin (10ug/mL, Corning, cat#354232). After coating, phNPCs were plated at a concentration of 400,000 cells per well. The cells were plated and fed with growth factors as described^22^. 24 hours after plating, the cells were half fed with the media and growth factors described^22^. 48 hours after plating, the cells were half-fed with the media and growth factors as described^22^, and with nucleic acid transfection enhancer (NATE) (1:100 final dilution; InvivoGen, cat#lyec-nate). 30 minutes after feeding, the cells were transfected with PJD_lib_minP_Luc plasmid via Lipofectamine Stem Transfection Reagent (Invitrogen, cat#STEM00001) according to the instructions, using Opti-Mem (Thermo Scientific, cat#31985070), 2.5ug of PJD_lib_minP_Luc plasmid, and 1ug of pUC19 plasmid (Thermo Scientific, cat#SD0061) per single 6-well. Co-transfecting with pUC19 plasmid increased transfection efficiency. 24 hours after transfection, the cells were half fed and then treated either with 7uL of 2.5uM CHIR or 7uL DMSO (vehicle). At exactly 48 hours after CHIR or vehicle treatment, RNA and DNA was extracted from the cells using the Quick-DNA/RNA Miniprep Plus Kit (Zymo, cat#D7003), with 600 µL of Shield Buffer and 600 µL of Lysis Buffer per well. In total, there were 16 vehicle replicates and 16 CHIR replicates, where one replicate was one well in a 6-well plate. For each 6-well plate, three wells were vehicle and three wells were CHIR treated.

### MPRA Library Preparation For Sequencing

Extracted RNA was reverse-transcribed into cDNA using SuperScript IV Reverse Transcriptase (Invitrogen, cat. no. 18090050) and a primer designed to bind downstream of the barcodes (Lib_Hand_RT). The resulting cDNA was then amplified with Lib_Hand and Lib_seq_Luc_R primers, utilizing NEBNext 2X Q5 Hifi HS Mastermix, and then purified using the Zymo DNA Clean and Concentrator kit-25 (Zymo, cat. no. D4033).

Additionally, 4 µL of DNA from each well was PCR amplified in four separate PCR reactions using Lib_Hand_RT and Lib_Seq_Luc_R primers with NEBNext 2X Q5 Hifi HS Mastermix, followed by purification with the Zymo Clean and Concentrator kit-25. P5 and P7 adapter sequences and i7 demultiplexing indices were then incorporated into both cDNA and DNA libraries through PCR using P5_seq_Luc_F and P7_Ind_#_Han primers with NEBNext 2X Q5 Hifi HS Mastermix.

The final PCR products were purified and size-selected using AMPure XP Beads (Beckman Coulter, cat. no. A63881) at 0.7X and 0.9X ratios to isolate a 273 bp DNA fragment. The libraries were then sequenced on a NovaSeq SP using a 35 × 8 × 0 cycle format with 20% PhiX and custom sequencing primers Index_primer and R1_primer (**Supplementary Table 8**).

### MPRA data processing

#### Sequencing alignment and Quantification of barcode counts

Adapter sequences were trimmed using cutadapt (v4.1)^56^ using default parameters. Trimmed sequences were then converted to reverse complement DNA sequences. We extracted only sequences with 20 bp length and perfectly matched to the BC information in the barcode-variant table which consists of the unique barcode-variant pairs.

In each replicate, we first filtered BCs if the DNA and RNA read counts had less than 5 read counts, and kept elements with at least 5 unique BC. Per each variant and replicate, log_2_(RNA/DNA) were fed to boxplot.stats() to detect BC outliers (**Supplementary Fig. 2**). After removing BC outliers, RNA and DNA counts were combined for each element. Among 16 replicates, those with lower correlation to other biological replicates (correlation coefficients to others = 0.3 - 0.7) were removed. Elements available at least 10 replicates were retained for analysis.

### MPRA regulatory effects

#### Regulatory activity

For each dataset (caQTL, ENIGMA-vehicle, ENIGMA-stimulation), α values (transcription rate adjusted by batch effects including sequencing depth) were estimated using *analyzeQuantification()* implemented in MPRAnalyze(v1.9.1)^57^ to assess regulatory activity of each element compared to the distribution of background (α of negative controls). *P*-values of median absolute deviation score (*P*-MAD score) were corrected for multiple comparisons by the Benjamini-Hochberg (BH) procedure [FDR]. Condition-dependent MPRA elements were identified among MPRA elements that were available in both conditions and were active in at least one of the conditions. Differential activity was estimated using *analyzeComparative()* followed by *testLtr()* in MPRAnalyze. For this analysis we excluded one replicate that had library size correction factors of DNA < 0.10 which led to the failure to execute *analyzeComparative()*. We set a statistical significant threshold at FDR < 0.1 to define “Active” elements or condition-dependent elements. In most subsequent analyses including enrichment analyses, each region was classified if either allele exhibited MPRA activity and as inactive if both alleles showed no detectable activity to ensure that each region was counted only once.

#### Allelic effect

We applied *mpralm()* (mpra v.1.16.0)^58^ followed by *topTable()* (limma v3.50.3)^59^ to detect allelic effects in MPRA active elements (MPRA element pairs with one of element was active in any conditions). Condition-dependent allelic effects were assessed with ∼genotype+condition+genotype*condition. Only MPRA element pairs that showed allelic differences (emVars) in at least one condition were used to test for condition-dependent effects. We set a statistical significance threshold at FDR < 0.1 to define emVars or condition-dependent emVars.

### Transposable elements

Transposable elements (TEs) overlapping MPRA elements were annotated using the RepeatMasker database obtained from the UCSC Table Browser. SINE, LINE, or LTR classes were extracted since those had at least 500 regions overlapped with MPRA elements used in this study (1,825, 1,367, 746, respectively). Alu elements, a subclass of SINEs, were further classified into three groups based on milliDiv score (<100, 100-150, >150)^35^ to interpret evolutionary ages.

#### Relative position of MPRA elements within the Alu sequence

To estimate the position of the MPRA element within the Alu element, the genomic location of each MPRA element was identified and overlapped with forward oriented Alu element that had length ≥ 250 bp. This position was used to estimate the approximate location of the A-Box (TGGCTCACGCC) and the B-Box (GTTCGAGAC), A_5_TACA_6_, and the poly-A Tail^33,34^. The relative MPRA element position within the Alu was determined by the distance from the Alu start position to the midpoint of the overlapping regions.

#### Enrichment analysis

Fisher’s exact tests were performed to assess the following enrichment of RE features in target groups against specified background.

1. Frequency of each TE overlaps (active vs inactive), (2) frequency of Alu overlaps (SINE-active vs SINE-inactive), (3) frequency of forwarded oriented Alu (Alu-active vs Alu-inactive), (4) frequency of A/B box followed by TTTT in 150bp, and (5) frequency of emVars in Alu elements (emVars vs MPRA active elements with non significant allelic effects).

When we tested enrichment in stimulation-enhancing MPRA activity, we used the number of RE in MPRA elements with non-significant activity differences between conditions as a reference while running Fisher’s exact test. We considered significant enrichment if FDR < 0.1.

### TF binding site motif enrichment test

To avoid testing redundant TF binding site [TFBS] motifs^60^, we used pre-clustered TF bind motifs available in JASPAR website (https://jaspar2020.genereg.net/static/clustering/2020/vertebrates/CORE/interactive_trees/JASPAR_2020_matrix_clustering_vertebrates_archive.zip]. Among 137 clusters, we extracted clusters in which at least one TF was expressed in phNPCs in either Vehicle or Wnt-stimulated condition^6^, resulting in 123 clusters. TFBSs within the 150 bp MPRA elements were predicted by FIMO (v5.5.2)^61^ with FDR < 0.1.

We evaluated (1) enrichment for any TF clusters overlaps, and (2) specific TF clusters using Fisher’s exact test.

### Enrichment analysis in various features for MPRA active elements

Since we tested MPRA activity per element not per variant, we extracted one element per variant with higher activity (i.e. smaller *P* value) for enrichment analyses unless specified. Those elements were then grouped as either active elements (FDR < 0.1) or inactive elements (FDR ≥ 0.1). In the inactive group, we ensured that both elements were tested and inactive. Two-tailed Fisher’s Exact tests were used to estimate the enrichment or depletion of MPRA active elements using MPRA inactive elements as a background set unless specified in the section below. A threshold of FDR < 0.1 was applied to identify statistically significant results.

### MPRA elements regulating their putative target genes

#### Endogenous gene regulatory information

Data were acquired from our previous study^6^. Protein-coding genes or lincRNAs expressed in unstimulated (vehicle) or CHIR stimulated phNPCs were extracted. Differentially expressed genes (DEGs) and differentially accessible chromatin peaks due to CHIR stimulation were defined if FDR < 0.1.

#### Mapping MPRA element to genes

We mapped MPRA elements to genes by overlapping either promoter regions defined as 2kb upstream of transcription start site (TSS) in Gencode v19 or chromatin interaction (Hi-C loops) from germinal zone (GZ) or cortical plate (CP). The proportion of MPRA elements linked to their putative target genes were compared between MPRA active elements vs MPRA inactive elements using Fisher’s exact test.

#### Directionality of condition-dependent effect

*binom.test()* was applied to test whether the percentage of concordant directionality of effect between MPRA activity and gene expression due to Wnt stimulation was higher than expected by chance.

### Prediction of TFs disrupted by emVars

motifbreakR (v2.8.0)^62^ were used to predict TFs disrupted by emVars on JASPAR2022 vertebrate from MotifDB with method=’ic’, threshold = 1e-4. We further extracted TFs predicted as strongly impacted (effect= “strong”) by motifBreakR and expressed in unstimulated and/or stimulated condition in phNPCs.

### CRISPRi experiments

gRNAs targeting a putative regulatory element near rs2802295 were designed using GT-Scan^63^ for CRISPRi experiments. An initial 4 targeting gRNAs positioned at roughly even spacing along the length of the regulatory element were selected in addition to 2 non-targeting gRNAs as controls. All gRNAs sequences tested are listed in **Supplementary Table 9** with the ones used for statistical analyses and high content screening in bold. gRNAs were cloned into pLV-U6-C-UbC-DsRed-P2A-Bsr (https://www.addgene.org/83919/) for gene expression experiments. Lenti-dCas9-KRAB-blast (https://www.addgene.org/89567/) vector was used for dCas9-KRAB expression. Lentiviruses for each plasmid were generated using the psPax2 (https://www.addgene.org/12260/) packaging plasmid and pMD2.G (https://www.addgene.org/12259/) envelope plasmid as described previously^5^. 293T cells were cultured directly in phNPC proliferation media for 48 hours after transfection using TransIT-LT1 Transfection Reagent (Mirus Bio). The viral supernatant was then harvested, purified by syringe filtering, and frozen at −80°C for direct use in subsequent experiments. Viral titers were calculated using the qPCR Lentivirus Titer Kit (Applied Biological Materials). A preliminary screen of the 4 gRNAs was performed to identify those with the highest potential for perturbing *FOXO3* expression. phNPCs were plated in 24 well plates at 80,000 cells per well. The following day, cells were infected with viruses for Lenti-dCas9-KRAB-blast and pLV-U6-C-UbC-DsRed-P2A-Bsr containing the targeting gRNAs at a calculated MOI of 2 for each virus. Plates were then spun at 37°C for 1 hour to increase transduction efficiency^64^. phNPCs were cultured for an additional 5 days after infection prior to RNA extraction.

RNA purification was performed using the Direct-Zol RNA Microprep Kit (Zymo Research) and cDNA from 200ng of purified RNA was generated using the iScript cDNA Synthesis Kit (Bio-Rad). Gene expression was then measured by qPCR using the SsoAdvanced Universal SYBR Green Supermix (Bio-Rad) and a QuantStudio 5 Real-Time PCR System (Thermo Fisher). *FOXO3* expression was normalized to the expression of the *ACTB* (Beta-actin) gene. Gene expression values for each targeting gRNA were compared to the average of the non-targeting controls. Primers used for qPCR are provided in **Supplementary Table 10.**

From our initial pilot screen, the 2 gRNAs out of 4 with the highest impact on gene expression were used for downstream experiments. qPCR experiments for each individual gRNA or a combination of 2 gRNAs were repeated for 5-6 unique phNPC donor lines with the average expression of 3 wells per donor per gRNA combination considered a biological replicate.

### Correlation between regional TF expression and GWAS SNP effects

Normalized gene expressions in fetal brains (8 post-conception weeks (pcw) to 24 pcw) were obtained from BrainSpan Atlas of the Developing Human Brain https://www.brainspan.org/static/download.html^65^. Gene expression was averaged for each cortical region defined by Allen Human Brain Atlas (AHBA) (**Supplementary Fig. 8**). GWAS Z scores were calculated using the effect size of the allele that matched the TF motif sequence divided by the standard error. Since brain regions used within the ENIGMA GWAS were based on Desikan-Killiany Atlas (DK-Atlas), we mapped 36 DK regions to 16 AHBA regions for comparison. Linear regression was used to assess correlation between GWAS Z score and gene expressions.

## Data availability

Data generated in this study are provided as supplementary tables, and raw data will be available upon publication.

## Code availability

All code used in analyses will be available before publication.

## Author Contributions

H.W. and J.L.S. conceived and supervised the study, and together with N.M., J.C.M., designed the study and data analysis strategy. J.C.M., O.K, and J.B. performed MPRA. N.M. conducted computational analyses. O.K. performed and analyzed CRISPRi experiment. B.D.L and J.M.V. provided Wnt dependent gene regulatory data. N.M., J.C.M., G.E.C., H.W. and J.L.S. interpreted the results. N.M., J.C.M, H.W., and J.L.S. wrote the initial draft of the manuscript. All co-authors provided critical feedback and approved the manuscript.

## Acknowledgements

This research was supported by the PsychENCODE consortium (R01MH122509, H.W. and J.L.S.). We acknowledge the technical support from the UNC High Throughput Sequencing Facility (HTSF; University Cancer Research Fund; Comprehensive Cancer Center Core Support grant, P30CA016086; UNC Center for Mental Health and Susceptibility grant, P30ES010126). We would like to acknowledge the use of bioRender (www.biorender.com) templates to make the cartoon in Fig. 1.

## Notes

### Competing Interest Statement

The authors have declared no competing interest.

### Summary of Updates

There was a misspelling in one author's name (Jessica C. McAfee). In addition small text edits were made.

