## Supplementary Information for "Massively parallel assessment of gene regulatory activity at human cortical structure associated variants"

### Supplementary Figures

#### Supplementary Figure 1| Reproducibility of caQTL

We compared the effect sizes of caQTLs identified in a population or primary human neural progenitors from two separate differentiations of largely the same donor lines derived from standard media[1] or media including a vehicle control[2]. Open circles indicate all caQTLs identified by previous study[2], and circles in red indicate caQTLs used in this MPRA study which were selected such that the caSNP was within the caPeak. The effect sizes are strongly correlated between studies (r = 0.87, $p < 2.2\times10^{-16}$ for all caQTLs and r = 0.98 < $2.2 \times 10^{-16}$ for the caQTLs used in the MPRA, respectively), indicating that the caQTLs were highly reproducible.

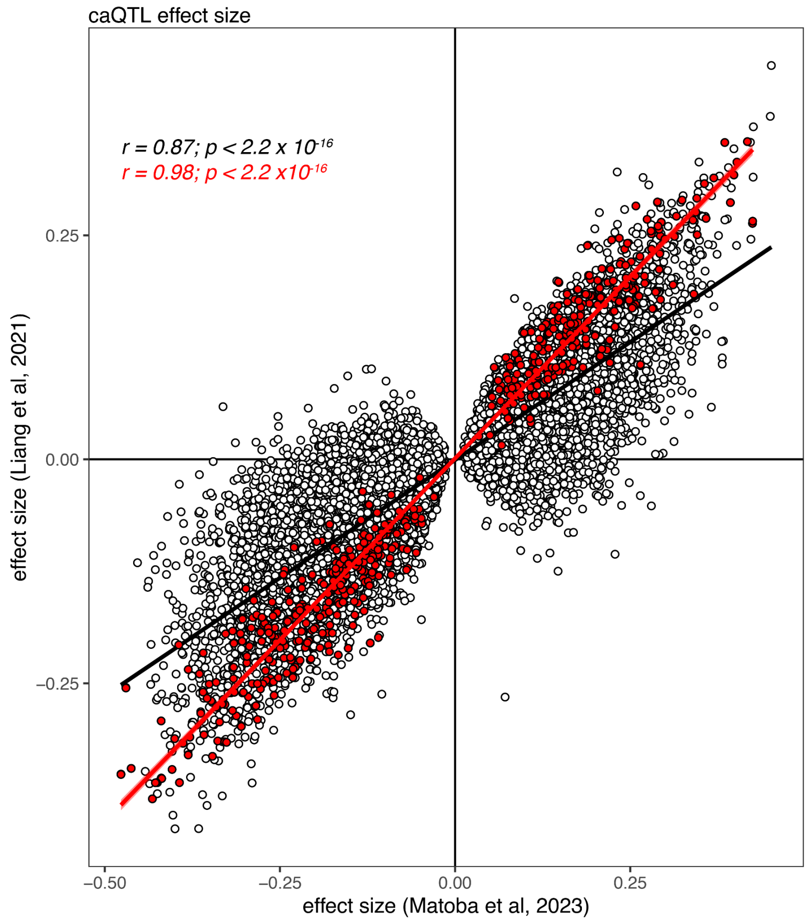

### Supplementary Figure 2| Examples of barcodes with outlier expression

Barcode outliers were identified by *boxplot.stats(log2(RNA/DNA))* function with the default setting in each variant within a replicate (**a**). We removed those outliers across replicates (**b**). Triangles in red represent outliers found in the replicate, and black represent outliers found in other replicate(s). Three examples from (1) positive control, (2) negative control, and (3) ENIGMA datasets are shown.

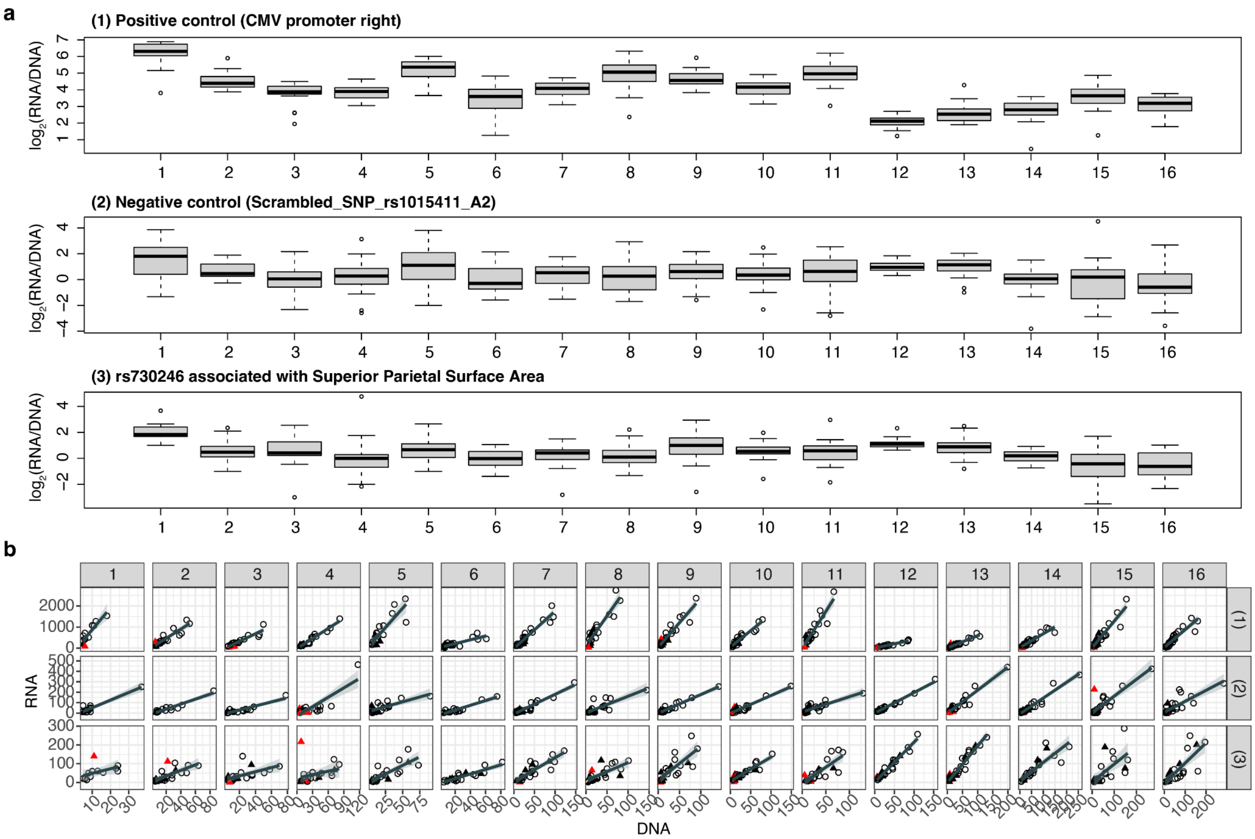

### Supplementary Figure 3| Barcode representation in MPRA element

For elements which survived quality control filtering (barcode representation > 5, present in replicates > 2, removal of barcode outliers based on activity), histograms show barcode count per element in vehicle condition (across element median = 28, (s.d.) = 12.2) (**a**) and stimulated condition (across element median = 29 (s.d. = 12.7) (**b**).

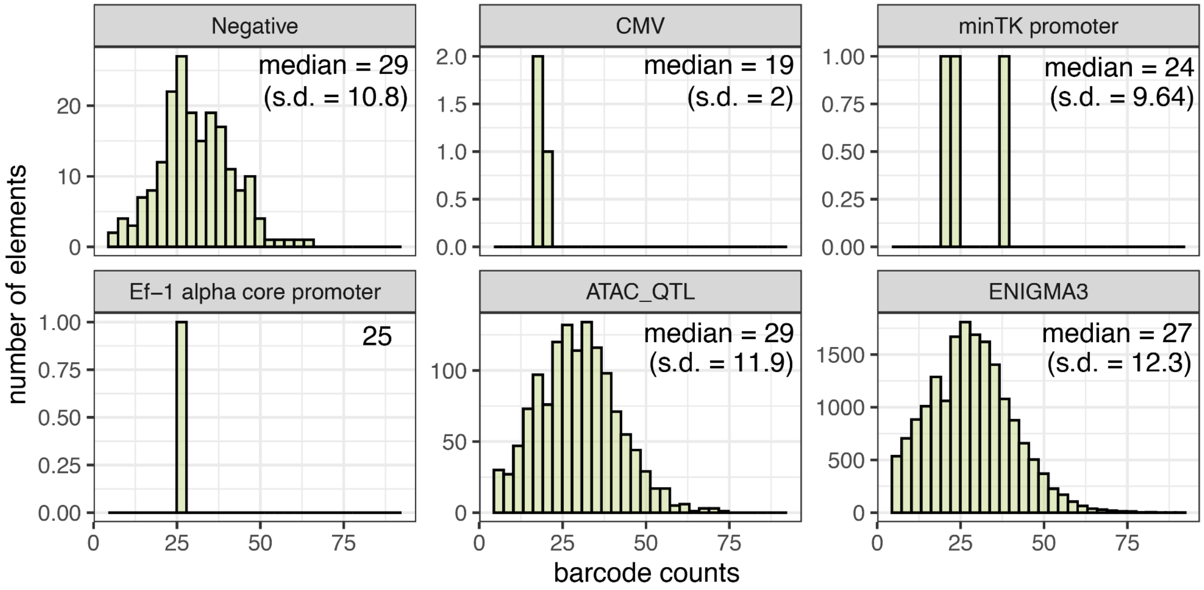

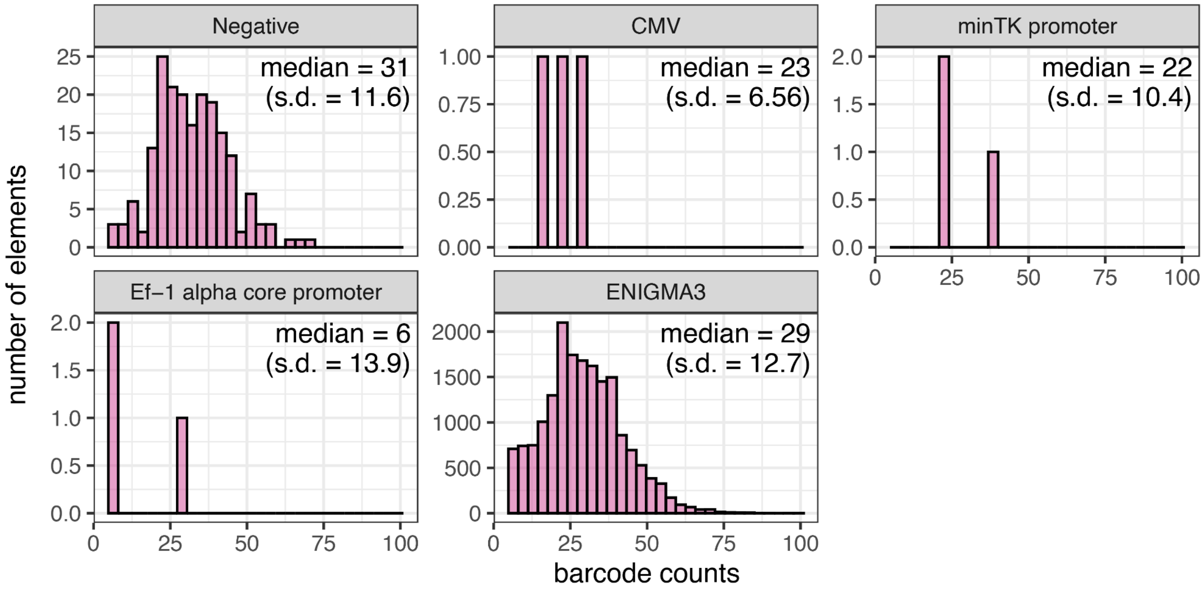

### Supplementary Figure 4| Positive controls show higher MPRA activity than negative controls

Scatter plot of DNA vs RNA counts of positive controls (colored in red) and negative controls (colored in blue) in replicate 1.

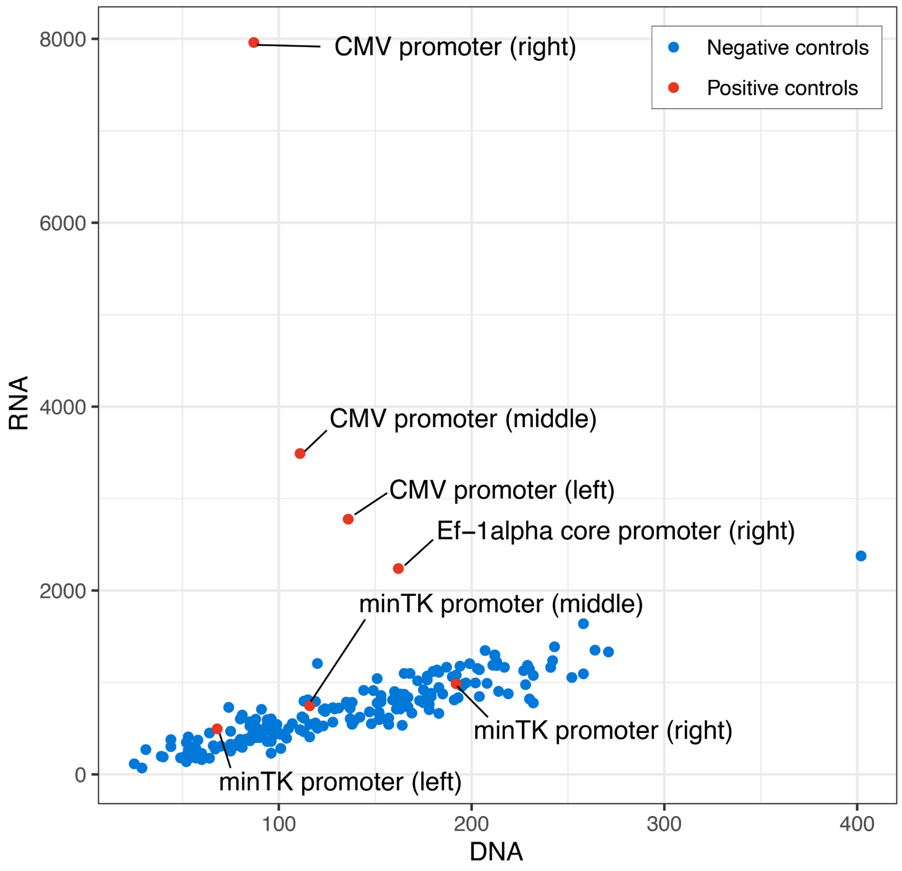

### Supplementary Figure 5| Limited concordance between exogenous multiplexed reporter and endogenous chromatin accessibility assays

**a.** Features correlated with caQTL MPRA active elements. Nominal p-values calculated by two-sided Wilcoxon Rank Sum test are given in the boxes, and the color corresponds to the signed $-\log_{10}$(p-value). The number of SNPs in high LD ($r^2 > 0.6$) with the caSNP within the 150 bp insert was significantly associated with MPRA activity, suggesting that haplotype effects might combinatorially increase MPRA activity, as has been found in previous work[3]. We also detected a nominally significant positive relationship between GC content of the insert and depth of ATAC coverage with the level of MPRA activity, suggesting that sequence biases or the proportion of cells with accessible regions lead to increased MPRA activity. The percentage of correlated peaks was also associated with decreased MPRA activity, suggesting that some regulatory elements cannot influence transcription on their own and instead must work together in networks to influence transcriptional activity. **b.** MPRA allelic effects ranked by effect size of the chromatin accessibility increasing allele compared with caQTL effect sizes. 6% of caQTL elements showed allelic effects in the same direction as MPRA (emVar(+), right) and 6% of caQTL elements showed allelic effects in the opposite direction as MPRA (emVar(-), left). **c.** No significant difference in caQTL effect size was detected between regions with MPRA allelic effects and regions without significant allelic effects. **d.** We detected no relationship between the allelic effect size in MPRA and the allelic effect size in caQTL. Colors indicate emVar(+) (red) or emVar(-) (blue) elements. **e-f**. Representative caQTLs for emVar(-) **(e)** and MPRA*+ **(f)** elements are shown. MPRA element (one per Locus) : $n_{\text{MPRA active}} = 30$, $n_{\text{MPRA inactive}} = 472$, $n_{\text{emVar++}} = 30$, $n_{\text{emVar-*-}} = 30$, $n_{\text{MPRAn.s\_allelic}} = 439$. $n_{\text{replicate}} = 14$.

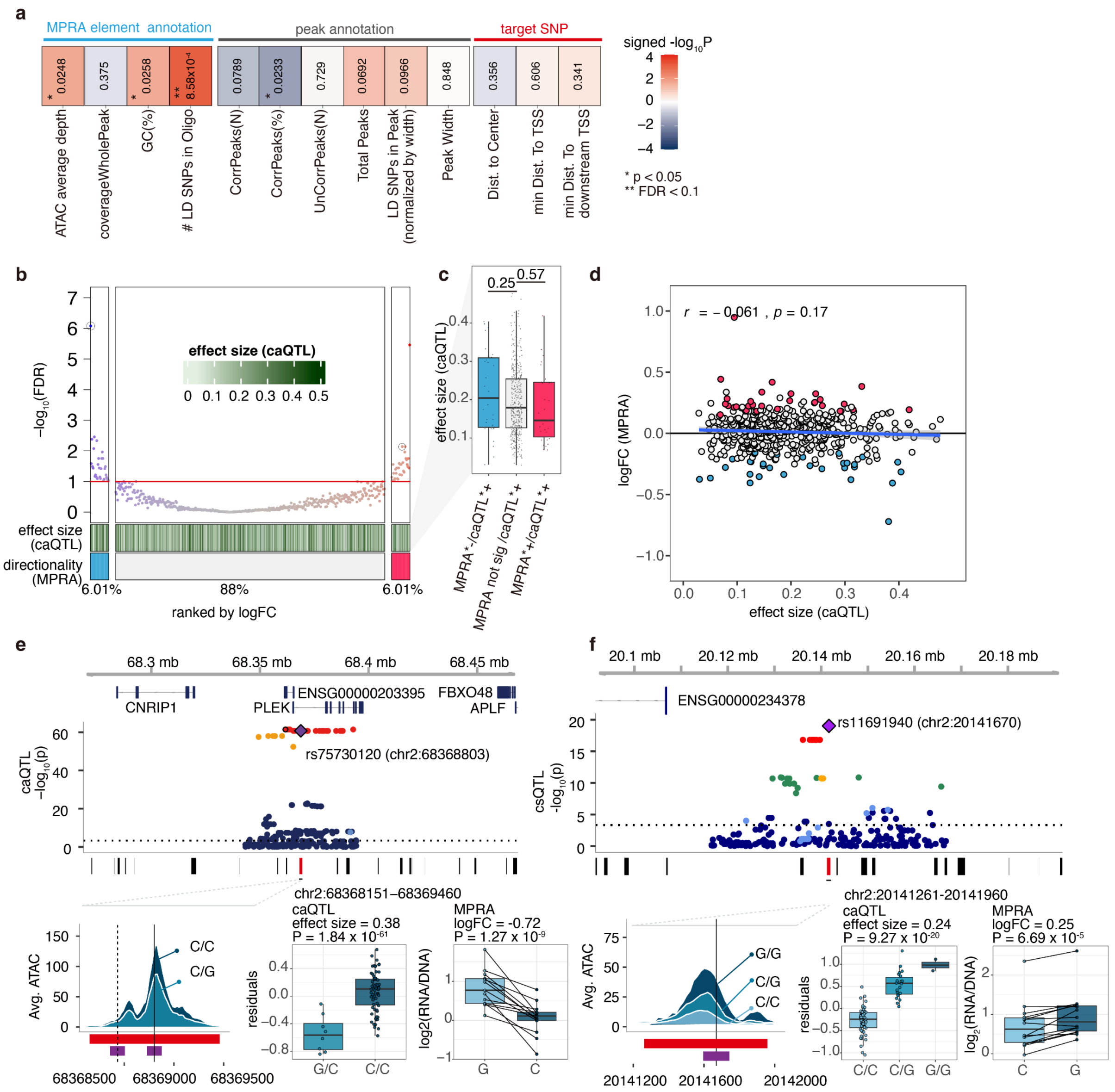
a
MPRA element annotation
peak annotation
target SNP
signed -log10P
* p < 0.05
** FDR < 0.1
ATAC average depth
coverageWholePeak
GC(%)
### LD SNPs in Oligo
CorrPeaks(N)
CorrPeaks(%)
UnCorrPeaks(N)
Total Peaks
LD SNPs in Peak (normalized by width)
Peak Width
Dist. to Center
min Dist. To TSS
min Dist. To downstream TSS
0.0248
0.375
0.0258
8.58x10-4
0.0789
0.0233
0.729
0.0692
0.0966
0.848
0.356
0.606
0.341
b
-log10(FDR)
effect size (caQTL)
effect size (caQTL)
directionality (MPRA)
6.01%
88%
6.01%
ranked by logFC
c
effect size (caQTL)
0.25
0.57
MPRA*-/caQTL*+
MPRA not sig /caQTL*+
MPRA*+/caQTL*+
d
r = −0.061 , p = 0.17
logFC (MPRA)
effect size (caQTL)
e
68.3 mb
68.35 mb
68.4 mb
68.45 mb
CNRIP1
ENSG00000203395
PLEK
FBXO48
APLF
caQTL -log10(p)
rs75730120 (chr2:68368803)
chr2:68368151–68369460
Avg. ATAC
C/C
C/G
68368500
68369000
68369500
caQTL
effect size = 0.38
P = 1.84 x 10-61
residuals
G/C
C/C
MPRA
logFC = -0.72
P = 1.27 x 10-9
log2(RNA/DNA)
G
C
f
20.1 mb
20.12 mb
20.14 mb
20.16 mb
20.18 mb
ENSG00000234378
rs11691940 (chr2:20141670)
csQTL -log10(p)
chr2:20141261-20141960
Avg. ATAC
G/G
C/G
C/C
20141200
20141600
20142000
caQTL
effect size = 0.24
P = 9.27 x 10-20
residuals
C/C
C/G
G/G
MPRA
logFC = 0.25
P = 6.69 x 10-5
log2(RNA/DNA)
C
G

#### Supplementary Figure 6| Overview of MPRA elements and cortical structure associated loci

**a.** UpsetR plot displays shared elements assessed for MPRA activity (top) and identified as MPRA active elements (bottom). Numbers in the barplot indicate element counts for either vehicle or stimulated condition (union) and each condition (vehicle, stimulation, respectively). Next to the UpsetR plot, the percentages of genomic features of 150 bp around the variant and allelic availability are shown. Genomic features are prioritized as follows: Exons > intron-exon boundaries > promoters > 1 to 5 kb from transcription start site [TSS] > introns > intergenic. **b**. Cortical-structure-associated loci ranked by the number of MPRA elements tested within the locus (top). Colors in heatmaps indicate phenotype-specific or shared locus where at least one MPRA variant is active stratified by condition. **c**. Histograms show the number of phenotype-specific loci (N = 55) and shared loci containing at least one MPRA active element. TH-SA: Thickness and surface area phenotypes; TH: Thickness; SA: Surface area.

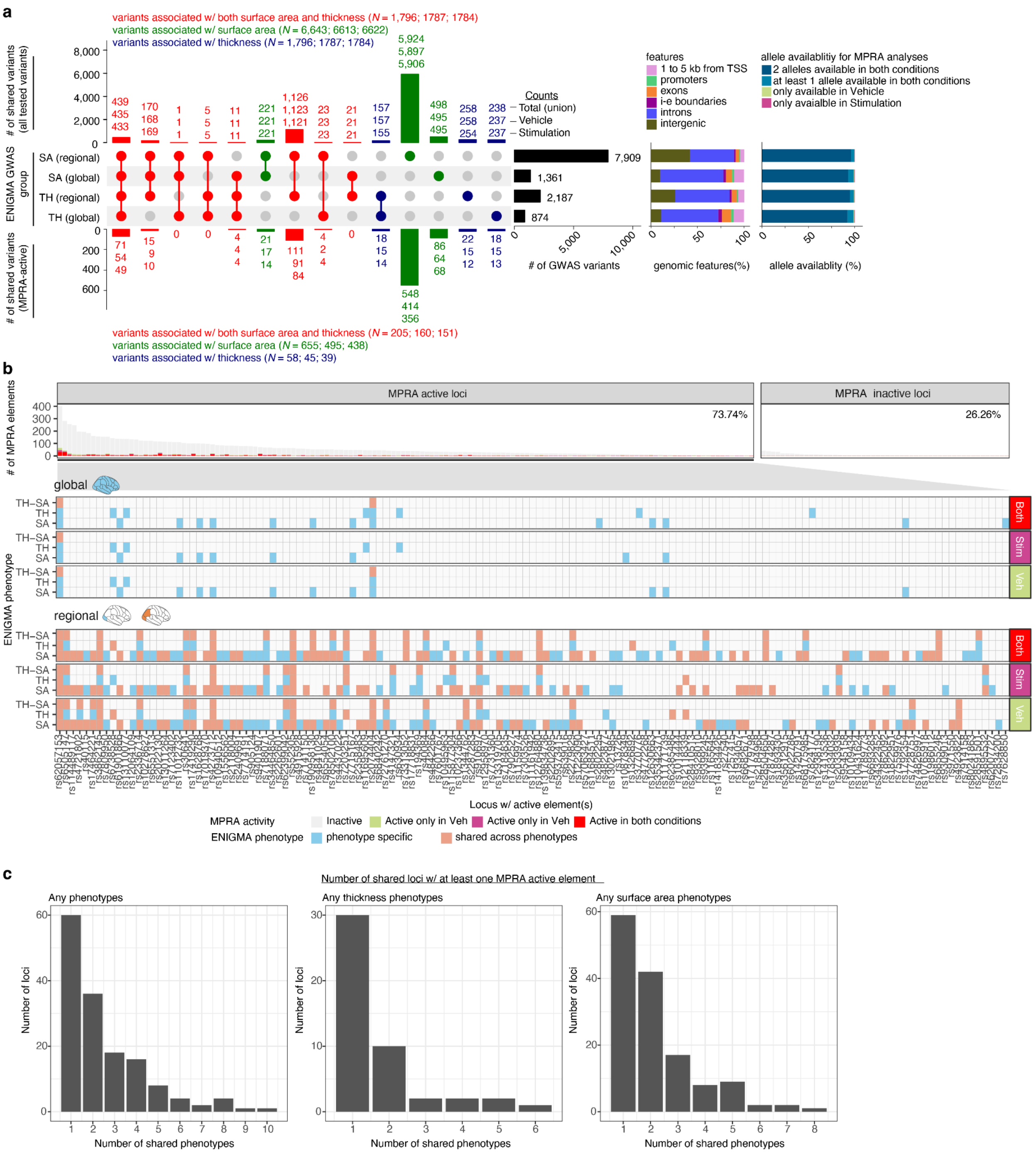
a
variants associated w/ both surface area and thickness (N = 1,796; 1787; 1784)
variants associated w/ surface area (N = 6,643; 6613; 6622)
variants associated w/ thickness (N = 1,796; 1787; 1784)
### of shared variants (all tested variants)
Counts
Total (union)
Vehicle
Stimulation
SA (regional)
SA (global)
TH (regional)
TH (global)
ENIGMA GWAS group
7,909
1,361
2,187
874
### of GWAS variants
features
1 to 5 kb from TSS
promoters
exons
i-e boundaries
introns
intergenic
genomic features(%)
allele availablitiy for MPRA analyses
2 alleles available in both conditions
at least 1 allele available in both conditions
only available in Vehicle
only avaialble in Stimulation
allele availablity (%)
### of shared variants (MPRA-active)
variants associated w/ both surface area and thickness (N = 205; 160; 151)
variants associated w/ surface area (N = 655; 495; 438)
variants associated w/ thickness (N = 58; 45; 39)
b
MPRA active loci
MPRA inactive loci
73.74%
26.26%
### of MPRA elements
global
regional
ENIGMA phenotype
TH–SA
TH
SA
Both
Stim
Veh
Locus w/ active element(s)
MPRA activity
Inactive
Active only in Veh
Active only in Veh
Active in both conditions
ENIGMA phenotype
phenotype specific
shared across phenotypes
c
Number of shared loci w/ at least one MPRA active element
Any phenotypes
Any thickness phenotypes
Any surface area phenotypes
Number of loci
Number of shared phenotypes

#### Supplementary Figure 7| Negative controls with a B-Box-like sequence show MPRA activity

Histograms of MPRA activity of negative controls quantified as transcription rate. Scrambled sequences generated from the rs73004638 region contain a B-Box-like sequence (GTTCGAGAC) (A1 = A, A2 = T at position 76), and show high transcription rate.

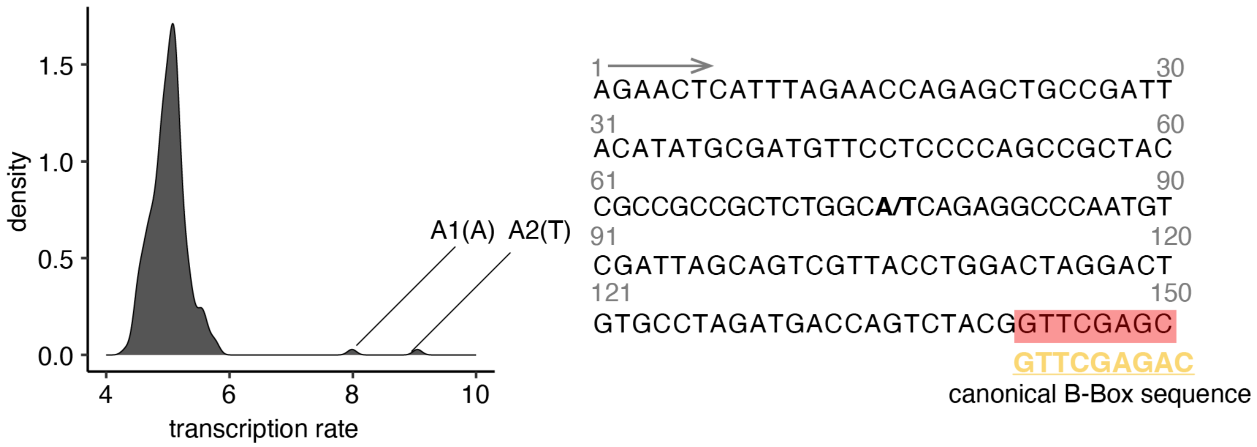

### Supplementary Figure 8| Projection of cortical regions between atlases

34 cortical regions from Desikan-Killiany Atlas (DK atlas) used in the ENIGMA dataset were mapped to 16 macroscale regions defined by The Allen Human Brain Atlas (AHBA).

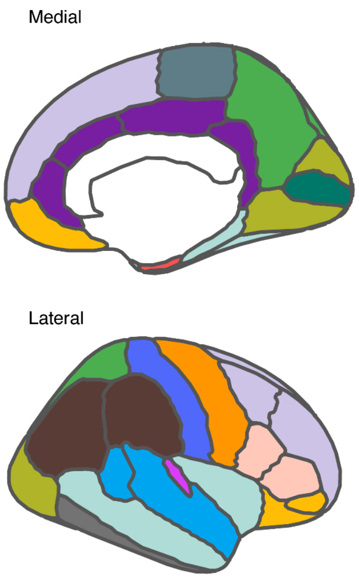

- Anterior (rostral) cingulate (medial prefrontal) cortex
  - Rostral anterior cingulate
  - Caudal anterior cingulate
  - Posterior cingulate
  - Isthmus cingulate
- Dorsolateral prefrontal cortex
  - Rostral middle frontal
  - Superior frontal
  - Caudal middle frontal
- Hippocampus (hippocampal formation)
  - Entorhinal
- Inferolateral temporal cortex (area TEv, area 20)
  - Inferior temporal
- Occipital neocortex
  - Lateral occipital
  - Lingual
  - Cuneus
- Orbital frontal cortex
  - Lateral orbitofrontal
  - Pars orbitalis
  - Frontal pole
  - Medial orbitofrontal
- Parietal neocortex
  - Superior parietal
  - Precuneus
- Posterior (caudal) superior temporal cortex (area 22c)
  - Banks of the superior temporal sulcus (bankssts)
  - Superior temporal
- Posteroventral (inferior) parietal cortex
  - Supramarginal
  - Inferior parietal
- Primary auditory cortex (core)
  - Transverse temporal
- Primary motor cortex (area M1, area 4)
  - Precentral
- Primary motor−sensory cortex (samples)
  - Paracentral
- Primary somatosensory cortex (area S1, areas 3,1,2)
  - Postcentral
- Primary visual cortex (striate cortex, area V1/17)
  - Pericalcarine
- Temporal neocortex
  - Fusiform
  - Parahippocampal
  - Temporal pole
  - Middle temporal
  - Insula
- Ventrolateral prefrontal cortex
  - Pars triangularis
  - Pars opercularis

#### Supplementary Figure 9| TFs associated with MPRA activity in the stimulated condition

**a**, Proportion of elements containing TF binding motifs stratified by MPRA activity. **b**, Enriched TF binding clusters. (**a**,**b**) Statistical significance of enrichment was assessed by a two-sided Fisher's exact test. **c**, The level of TF motif disruption (x-axis) is related to MPRA allelic differences (y-axis). P-values were estimated by Pearson's correlation test.

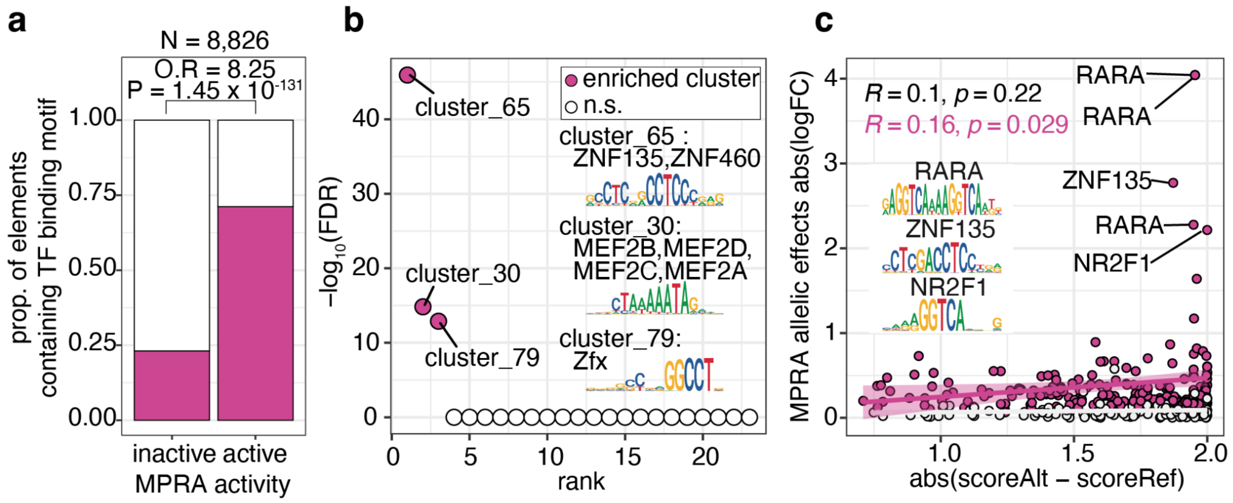

### Supplementary Tables

Supplementary Tables 2 and 3 are provided in Excel files.

#### Supplementary Table 1| Summary of the tested elements after QC

| | | caQTL | | ENIGMA | | | | | |
|---|---|---|---|---|---|---|---|---|---|
| | | all | paired | all | | | paired | | |
| | Condition | Veh | Veh | Veh | Stim | Union | Veh | Stim | Union |
| A | Elements | 1,316 | 1,310 | 17,837 | 17,848 | 17,980 | 17,570 | 17,592 | 17,756 |
| B | Variants | 661 | 655 | 9,052 | 9,052 | 9,092 | 8,785 | 8,796 | 8,878 |
| C | Locus | 502 | 499 | 198 | 198 | 198 | 198 | 198 | 198 |
| D | Active elements | 37 | 34 | 988 | 927 | 1,258 | 949 | 897 | 1,202 |
| E | Active variants | 33 | 30 | 700 | 628 | 918 | 661 | 598 | 863 |
| F | Active locus | 30 | 28 | 138 | 133 | 150 | 135 | 131 | 146 |
| G | % active elements (D/A) | 2.81 | 2.60 | 5.54 | 5.19 | 7.00 | 5.40 | 5.10 | 6.77 |
| H | % active variants (E/B) | 4.99 | 4.58 | 7.73 | 6.94 | 10.10 | 7.52 | 6.80 | 9.72 |
| I | % active Locus (F/C) | 5.98 | 5.61 | 69.70 | 67.17 | 75.76 | 68.18 | 66.16 | 73.74 |
| J | emVars | | 13 | | | | 386 | 333 | 536 |
| K | Locus with emVar(s) | | 12 | | | | 107 | 107 | 126 |
| L | % emVars (J/E) | | 43.33 | | | | 58.40 | 55.69 | 62.11 |
| M | % locus with emVar(s) (K/F) | | 42.8 | | | | 79.26 | 81.68 | 86.30 |

#### Supplementary Table 4| Summary of correlation between TF expression level across brain area and GWAS effect size (detailed statistics are provided in Supplementary Table 5)

| GWAS measurement | SNP disrupts motifs | Number of TFs | TFs (coefficient; BH-adjusted P) |
|---|---|---|---|
| Surface area | rs12046466 | 2 | RARG (1.34; 0.00611)<br>SREBF1 (1.34; 0.0465) |
| Surface area | rs1910326 | 4 | EOMES (−1.59; 0.0688)<br>SNAI3 (1.61; 0.0637)<br>TBX15 (−1.41; 0.0061)<br>TWIST1 (1.97; 0.0061) |
| Surface area | rs2395882 | 1 | ZNF85 (−0.845; 0.0686) |
| Surface area | rs28735393 | 3 | CREB1 (−1.25; 0.0686)<br>RARG (−1.66; 0.000483)<br>RORB (1.34; 0.0686) |
| Thickness | rs11887401 | 1 | FOSL2 (−1.16; 0.0275) |
| Thickness | rs28495472 | 1 | SREBF2 (1.02; 0.0688) |
| Thickness | rs62064600 | 1 | SREBF2 (1.29; 0.0768) |

#### Supplementary Table 5| Correlation between TF expression level across cortical areas and GWAS effect size (Related to Supplementary Table 4)

| GWAS measurement | SNP | TF | coef | SE | t value | p_value | BH-adjusted-P |
|---|---|---|---|---|---|---|---|
| Surface area | rs12046466 | RARG | 1.34 | 0.277 | 4.84 | $3.13 \times 10^{-5}$ | 0.0061 |
| Surface area | rs12046466 | SREBF1 | 1.34 | 0.339 | 3.96 | $3.97 \times 10^{-4}$ | 0.0465 |
| Surface area | rs1910326 | EOMES | -1.59 | 0.451 | -3.53 | $1.29 \times 10^{-5}$ | 0.0688 |
| Surface area | rs1910326 | SNAI3 | 1.61 | 0.425 | 3.78 | $6.52 \times 10^{-4}$ | 0.0637 |
| Surface area | rs1910326 | TBX15 | -1.41 | 0.422 | -3.33 | $2.18 \times 10^{-3}$ | 0.0984 |
| Surface area | rs1910326 | TWIST1 | 1.97 | 0.395 | 4.98 | $2.14 \times 10^{-5}$ | 0.0061 |
| Surface area | rs2395882 | ZNF85 | -0.85 | 0.233 | -3.62 | $9.96 \times 10^{-4}$ | 0.0686 |
| Surface area | rs28735393 | CREB1 | -1.25 | 0.345 | -3.63 | $9.79 \times 10^{-4}$ | 0.0686 |
| Surface area | rs28735393 | RARG | -1.66 | 0.272 | -6.09 | $8.25 \times 10^{-7}$ | 0.00048 |
| Surface area | rs28735393 | RORB | 1.34 | 0.371 | 3.60 | $1.05 \times 10^{-3}$ | 0.0686 |
| Thickness | rs11887401 | FOSL2 | -1.16 | 0.275 | -4.22 | $1.88 \times 10^{-4}$ | 0.0275 |
| Thickness | rs28495472 | SREBF2 | 1.02 | 0.29 | 3.53 | $1.29 \times 10^{-3}$ | 0.0888 |
| Thickness | rs62064600 | SREBF2 | 1.29 | 0.374 | 3.46 | $1.57 \times 10^{-3}$ | 0.0768 |

### Supplementary Table 6| Primer information for oligo library

| Primer | Sequence |
|---|---|
| MPRA-chipprimer-F | TATGCTGGTATACGCCTACA |
| MPRA-chipprimer-R | CCGTCACTAACTAACAGTGG |
| MPRA-BC_Primer_R | /5BiosG/AGTCGACTAGTNNNNNNNNNNNNNNNNNNNNTCTAGACCGTCACTAACTAACAGTGG |

### Supplementary Table 7| Primer information for barcode mapping

| Primer | Sequence |
|---|---|
| Bcmap_P5_AAV_R | AATGATACGGCGACCACCGAGATCTACACCGATAAGCTTGATATCGAATTCCC |
| Bcmap_P7_AAV_F_index | CAAGCAGAAGACGGCATACGAGATNNNNNNATTATCACTAGGGGTTCCTGC |
| BCmap_R1Seq_AAV_R | CGATAAGCTTGATATCGAATTCCCTAATGCACTAGT |
| BCmap_R2Seq_AAV_F | ATTATCACTAGGGGTTCCTGCGGCCGCA |
| BC_Map_R1seq | GTAACCACCCTGATCGACGGGGAGTGTACTAGT |

### Supplementary Table 8| Primer information for MPRA sequencing

| Primer | Sequence |
|---|---|
| Lib_Hand_RT | ATGCTCTTCCCAACTGCCGACGGGGAGTGTACTAGT |
| Lib_Hand | TGCTCTTCCCAACTGCCGA |
| Lib_seq_Luc_R | TACAACCGCCAAGAAGCTGC |
| P5_seq_Luc_F | AATGATACGGCGACCACCGAGATCTACACTACAACCGCCAAGAAGCTGC |
| P7_Ind_#_Han | CAAGCAGAAGACGGCATACGAGATNNNNNNNNGCGTGCTCTACGACTATGCTCTTCCCAACTGCCGA |
| Index_primer | TCGGCAGTTGGGAAGAGCATAGTCGTAGAGCACGC |
| R1_primer | CCAAGAAGGGCGGCAAGATCGCCGTGTAATAATTCTAGA |

#### Supplementary Table 9| CRISPRi screening gRNA sequences

| Name | Position | Strand | Sequence (5 ' - 3') |
|---|---|---|---|
| Control-1 | NA | NA | GAACCTCCCCGAATATCTGG |
| Control-2 | NA | NA | GTATTACTGATATTGGTGGG |
| FOXO3-RE2-76 | chr6:108,927,356 | + | GGAATTCTGTACAGTATGCTGGG |
| FOXO3-RE2-259 | chr6:108,927,539 | + | CCTTCCCAAGGGCGGTTTGTTGG |
| **FOXO3-RE2-263** | **chr6:108,927,543** | **-** | **CTTGCCAACAAACCGCCCTTGGG** |
| **FOXO3-RE2-474** | **chr6:108,927,754** | **+** | **TATGATTAAGGCTTATCTACAGG** |

#### Supplementary Table 10| qPCR primers for CRISPRi screening

| Target gene | Strand | Sequence (5 ' - 3') |
|---|---|---|
| *ACTB* | + | CATGTACGTTGCTATCCAGGC |
| *ACTB* | - | CTCCTTAATGTCACGCACGAT |
| *FOXO3* | + | CGGACAAACGGCTCACTCT |
| *FOXO3* | - | GGACCCGCATGAATCGACTAT |
